# *CHD8* haploinsufficiency alters the developmental trajectories of human excitatory and inhibitory neurons linking autism phenotypes with transient cellular defects

**DOI:** 10.1101/2020.11.26.399469

**Authors:** Carlo Emanuele Villa, Cristina Cheroni, Alejandro López-Tóbon, Christoph P. Dotter, Bárbara Oliveira, Roberto Sacco, Aysan Cerag Yahya, Jasmin Morandell, Michele Gabriele, Christoph Sommer, Mariano Gabitto, Giuseppe Testa, Gaia Novarino

## Abstract

Chromodomain helicase DNA-binding 8 (*CHD8*) is one of the most frequently mutated genes causative of autism spectrum disorder (ASD). While its phenotypic spectrum often encompasses macrocephaly and hence implicates cortical abnormalities in this form of ASD, the neurodevelopmental impact of human *CHD8* haploinsufficiency remains unexplored. Here we combined human cerebral organoids and single cell transcriptomics to define the effect of ASD-linked *CHD8* mutations on human cortical development. We found that C*HD8* haploinsufficiency causes a major disruption of neurodevelopmental trajectories with an accelerated generation of inhibitory neurons and a delayed production of excitatory neurons alongside the ensuing protraction of the proliferation phase. This imbalance leads to a significant enlargement of cerebral organoids aligned to the macrocephaly observed in patients with *CHD8* mutations. By adopting an isogenic design of patient-specific mutations and mosaic cerebral organoids, we define genotype-phenotype relationships and uncover their cell-autonomous nature. Finally, our results assign different *CHD8*-dependent molecular defects to particular cell types, pointing to an abnormal and extended program of proliferation and alternative splicing specifically affected in, respectively, the radial glial and immature neuronal compartments. By identifying temporally restricted cell-type specific effects of human *CHD8* mutations, our study uncovers developmental alterations as reproducible endophenotypes for neurodevelopmental disease modelling.

## Introduction

Autism spectrum disorders (ASDs) are a group of heterogeneous conditions characterized by difficulties in establishing social contacts and the manifestation of repetitive behaviors. Currently, there are no cures for the core symptoms of ASD, reflecting a lack of understanding of the major events leading to this class of neuropsychiatric disorders. Investigating how mutations in high-ranking risk genes lead to ASD remains a challenge. Functional genomic studies and single cell RNA sequencing of postmortem brain samples obtained from autistic patients have highlighted alterations of transcription in pyramidal cells and local interneurons at early to mid-fetal developmental stages^1–3^, however evidence from tractable experimental models of human neurodevelopment is still lacking. One of the major limitations is the inaccessibility of the affected tissue during development, when the bases of these disorders are eventually laid out. The possibility to generate human cerebral organoids from embryonic or pluripotent stem cells (hESC and iPSC) represents a new opportunity to study the cellular and molecular consequences of ASD-associated mutations. Cerebral organoids represent the closest *in vitro* model of a human cortex at early embryonic stages^4,5^, thus providing a unique tool to unravel the impact of gene impairment during the ASD vulnerable time windows^3^.

Genes coding for chromatin-remodelers are frequently mutated in ASD patients^6,7^ and among them chromodomain helicase DNA binding protein 8 (*CHD8*) is one of the most frequently mutated and most penetrant^8–10^. *CHD8*, initially identified in a screen for interactors within the canonical Wnt/beta-catenin pathway^11^, is part of the chromodomain-helicase-DNA binding protein family, characterized by a SFN2-like ATPase and two chromo-domains. The majority of the ASD-associated *CHD8 de novo* mutations lead to loss-of-function (LoF) and results in gene haploinsufficiency^8–10,12^. Patients with such mutations, in addition to autism-relevant behaviors, present with gastrointestinal complaints, intellectual disability and macrocephaly, linking *CHD8* haploinsufficiency to abnormal cortical development^8,12^.

Previous studies have explored the role of *CHD8* haploinsufficiency in brain development and ASD manifestations by employing animal models^13–16^ or iPSC-derived models profiled at a single time point and in bulk^17^, thus precluding the investigation of neurodevelopmental trajectories. Importantly, while in humans *CHD8* LoF mutations lead to a number of severe neurological problems, in mouse heterozygous mutations are associated with rather mild neurological abnormalities^14–16^. In addition, while *Chd8* haploinsufficient mice show a slight but significant increase of brain size, *in utero* shRNA-mediated *Chd8* knockdown results in reduced neural progenitor cell proliferation and premature neuronal differentiation^13^, a phenotype usually associated with reduced brain size. Thus, overall, the cellular and molecular mechanisms underlying *CHD8*-associated phenotypes remain still elusive, and conflicting results suggest possible technique- and species-specific issues, making it difficult to find a consensus on the role of *CHD8* in the pathogenesis of ASDs. Here we employed a cerebral organoid model system to determine the effects of *CHD8* haploinsufficiency on human cortical development. We found that *CHD8* haploinsufficiency leads to a significant enlargement of cerebral organoids with associated features of abnormal cortical development. Sequencing of RNA from 75060 single cells from control and *CHD8* mutant organoids at different developmental stages revealed a disrupted temporal differentiation dynamic between excitatory and inhibitory neurons, including a longer phase of excitatory neuron progenitor proliferation, in parallel to an accelerated production of inhibitory neurons. At the molecular level our analysis uncovers cell-type specific phenotypes that underlie differentially impacted developmental trajectories and point to temporally discrete mechanisms.

## Results

### Generation and characterization of human embryonic stem cell-derived cerebral organoids

We and others recently proposed that the combination of disease-specific organoid models and single cell omic resolution are materializing the vision of interceptive medicine, whereby deviations from development and homeostasis are intercepted and recast as problems of cellular dynamics within tissues^18^. A prerequisite for this approach is the reproducibility of organoid generation in longitudinal designs, both within and across batches, so as to enable robust phenotyping. Reproducibility issues have been widely discussed in the field and have led to the proposition of a number of alternative protocols for the generation of cerebral organoids, each aiming to improve in- and inter-batch reproducibility and/or specifically driving the generation of defined neuronal cell types^19–22^. In order to establish reliable models of *CHD8* haploinsufficiency, we considered that reproducibility problems may likely arise during the ectoderm and neuroectoderm specification in embryoid bodies (EBs). Indeed, suboptimal timing in the induction of the neuroectoderm radial organization at this stage compromises the subsequent formation of the neuroepithelium in stereotypical rosette-like structures budding from the main EB body^23^, likely indicating unstable early ectoderm formation in the EB right after hESC seeding. Building on other successful methods^4^, we reasoned that optimizing culture conditions during the hESC seeding step might be key to further increase the robustness of EB formation. Specifically, we took inspiration from work^5^ showing strong positive correlation between human fetal and organoid cell types and introduced slight variations to the initial culturing steps of the original protocol (see Material and Methods). Indeed, in our culture conditions (Fig. 1a), we observed a high degree of reproducibility in the quality of EB ectoderm and neuroectoderm differentiation (Extended Data Fig 1a), which, as expected, resulted in a striking high number of successfully differentiated cerebral organoids per number of matrigel embedded EBs (>90%). We verified by immunostaining that the reproducibility observed at the macroscopic level was matched by a reproducible cell composition in the cortical structures (Extended Data Fig. 1b, c). Similar to what was previously shown^23^, control cerebral organoids displayed the progressive features of cortical development starting with the formation of ventricular zone-like structures (VZ), the appearance of subventricular-like areas (SVZ) and the development of organized neuronal cell layers (Extended Data Fig. 1d). In agreement with the developing human cortical VZ, VZ-like structures of cortical organoids at day 10 are constituted almost exclusively by Sox2-positive radial glia cells (Extended Data Fig. 1e). As time progresses an increasing proportion of intermediate progenitors (TBR2 positive) and lower layer (CTIP2 positive) and upper layer (SATB2 positive) neurons can be observed, indicating the formation of structures reminiscent of the cortical-plate and a layered structure (Extended Data Fig. 1f). As development progresses, the size of the organoids increases accordingly (Fig. 1a). Finally, we checked *CHD8* expression at multiple stages and found its mRNA levels to be highest in 36-day old cerebral organoids (Fig. 1b), suggesting that *CHD8* may play essential roles around that developmental stage.

**Figure 1.**
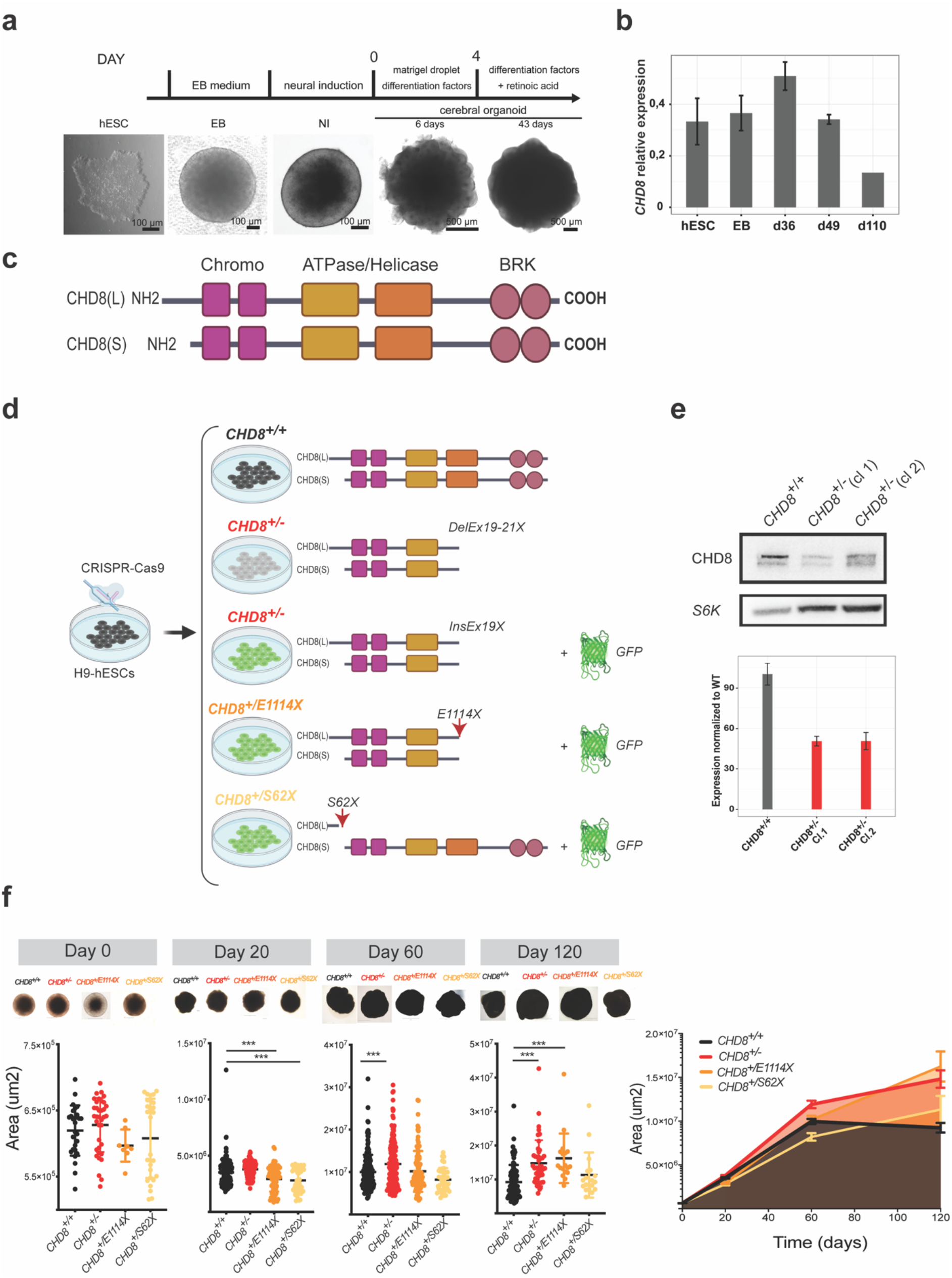
Increased cerebral organoid size linked to CHD8 heterozygous mutations. (**a**) Outline for the generation of cerebral organoids from hESCs. (**b**) Results of quantitative RT-PCR were normalized to *TBP* and relative expression values are shown for each time point. *CHD8* is expressed during all stages of cerebral organoidogenesis. Results are presented as mean ± s.e.m. (**c**) Schematics of the human long (L) and short (S) CHD8 isoforms. (**d**) Schematics illustrating the CRISPR/Cas9-mediated generation of the hESC lines employed in this study, the resulting genotypes and predicted proteins. DelEx19-21 X: deletion of part of exon 19, whole exon 20 and part of exon 21 and premature STOP codon; InsEx19X: eGFP was inserted inside the exon 19 with the concomitant introduction of a premature STOP codon; E1114X: the codon encoding for the glutamate in position p.1114 was mutated to generate a STOP codon; S62X: the serine in position 62 was mutated to a STOP codon affecting specifically only the long isoform. (**e**) Western blot of *CHD8*^+/+^ and *CHD8*^+/−^ hESCs revealing reduction in *CHD8* protein and mRNA levels upon deletion of exon 20, normalized to S6K. Expression levels are shown relative to *CHD8*^+/+^. Error bars display technical variation from triplicates. (**f**) Representative pictures and area of the cerebral organoids of the four different genotypes at day 0, 20, 60 and 120. For this analysis *CHD8*^+/−^ and *CHD8*^+/−;GFP^ organoids were pulled together. Day 0 n(*CHD8*^+/+^) =27; n(*CHD8*^+/+^) =36; n(*CHD8*^+/E1114X^) =11; n(*CHD8*^+/S72X^) =27. Day 20 n(*CHD8*^+/+^) =196; n(*CHD8*^+/+^) =119; n(*CHD8*^+/E1114X^) =80; n(*CHD8*^+/S72X^) =45. Day 60 n(*CHD8*^+/+^) =202; n(*CHD8*^+/+^) =163; n(*CHD8*^+/E1114X^) =106; n(*CHD8*^+/S72X^) =41. Day 120 n(*CHD8*^+/+^) =85; n(*CHD8*^+/+^) =46; n(*CHD8*^+/E1114X^) =18; n(*CHD8*^+/S72X^) =18. N indicates number of organoids. ***P< 0.001, Ordinary One-way ANOVA followed by Dunnett’s multiple comparisons test.

### Generation of isogenic *CHD8* haploinsufficient hESC lines

We built on our previous transcriptomics-based disease modelling benchmark^24^ and adopted an isogenic experimental design introducing *CHD8* mutations in an isogenic human embryonic stem cell (hESC) line employing the Clustered Regularly Interspaced Short Palindromic Repeats (CRISPR)/Cas9 genome engineering technology^25,26^.

In humans the *CHD8* gene has a short and a long isoform, with the former lacking the first 279 N-terminal amino acids (Fig. 1c). Both isoforms however display one helicase and two chromatin binding domains (Fig. 1c). To study the effect of *CHD8* LoF mutations we generated two separate hESC lines (*CHD8^+/−^* and *CHD8^+/−;GFP^*) carrying a deletion in the genomic portion of *CHD8* encoding for the C-terminal helicase domain, as well as two cell lines carrying patient specific mutations resulting in premature stop codons (i.e. S62X and E1114X) (Fig. 1d). Importantly, while both patient mutations have been linked to the autism core symptoms, the S62X patient does not present with macrocephaly and intellectual disability^8,12^, indicating that the S62X mutation results in only a subset of the *CHD8* clinical phenotypes. In all but one case (i.e. *CHD8^+/−^*) we coupled *CHD8* mutations with the expression of enhanced green fluorescence protein (eGFP) (Fig. 1d). This approach gave us the possibility to easily track mutant cells and perform experiments comparing wild type and mutant cells side by side. Human ES cells carrying the selected *CHD8* mutations on a single allele (i.e. *CHD8^+/−^*, *CHD8^+/−;GFP^*, *CHD8^+/E1114X;GFP^*, *CHD8^+/S62X;GFP^*) were isolated, expanded and analyzed for potential off target effects (Extended Data Fig. 2 and Material and Methods). Next, we tested the effect of the mutations on CHD8 protein levels by immunoblotting and confirmed that all the mutants show reduced levels of CHD8 protein (Fig. 1e and Extended Data Fig. 3).

### *CHD8* haploinsufficiency leads to a macrocephaly-like phenotype in human cerebral organoids

We employed the *CHD8* mutant and control hESC lines to generate human cerebral organoids. Similar to the mutant hESC lines, *CHD8* mutant organoids show a 50% reduction of CHD8 protein levels when compared with *CHD8^+/+^* samples (Extended Data Fig.3). At very early stages (matrigel embedding, day 0) control (*CHD8^+/+^*) and mutant (*CHD8^+/−^*, *CHD8^+/E1114X^*, *CHD8^+/S62X^*) cerebral organoids are indistinguishable, with equal appearance and size as well as comparable neuroectoderm thickness (Fig. 1f). Surprisingly, 20 days after matrigel embedding *CHD8* mutant organoids are however either equally large or slightly smaller than their control counterpart (Fig. 1f). This observation was in contrast to what was expected given that LoF mutations in *CHD8* are associated with macrocephaly. We thus let the cerebral organoids grow longer and found that by day 60 *CHD8^+/−^* and *CHD8^+/E1114X^* organoids are comparable or significantly larger than wild type samples (Fig. 1f). This trend becomes even more obvious by day 120, when *CHD8^+/−^* and *CHD8^+/^*^E1114X^, but not *CHD8^+/S62X^*, cerebral organoids are about 50% larger than controls (Fig. 1f). Thus, *CHD8* haploinsufficiency results in enlargement of human cortical organoids between day 20 and day 120 (Fig. 1f). Importantly, S62X has hardly any effect on organoid size by day 120, confirming that this mutation is not associated with macrocephaly and underscoring the sensitivity of our organoid system in capturing pathophysiological readouts of patient-specific mutations.

### Effects of *CHD8* haploinsufficiency at single cell resolution

To define at high resolution the cellular mechanisms underlying *CHD8* haploinsufficiency phenotypes we comprehensively analyzed control and mutant organoids by droplet-based single cell RNA sequencing (scRNAseq). Specifically, we profiled gene expression of 75060 cells obtained from a longitudinal cohort amounting to a total of 63 control and mutant cerebral organoids (Fig 2a, e) at three different developmental stages (i.e. day 20, day 60 and day 120) (Fig. 2a). After data integration (see Material and Methods), application of Leiden, a community detection algorithm optimized to identify communities guaranteed to be connected and hence to reliably score cell population based phenotypes^27^, yielded 10 different cell clusters. To characterize the resulting cellular clusters, we next adopted an integrative annotation strategy combining: i) supervised plotting of known cell identity markers (Extended Data Figure 4a); ii) cluster-wise overlap of cluster identity markers in organoids vs human fetal cortex^28^ and iii) integration, in the same dimensionality reduction space (UMAP), of the cell populations from our cerebral organoid dataset with the ones from a large public dataset of fetal human primary cells spanning several developmental stages^29^ (Extended Data Figure 4c,d). This annotation strategy led to the discrimination of 10 cell populations including three clusters of radial glia cells (RGCs) comprising different cell cycle phases (Fig. 2a-d, RG1, RG2 and RG3), intermediate progenitors (IP), interneuron progenitors (IN_IP), interneurons (IN), early excitatory neurons (ENE), excitatory neurons of upper (EN1) and lower (EN2) layers along with a minor number (2,9%) of unidentified cells (Fig. 2a, Extended Data Fig. 4). To determine the evolution of the system in terms of temporal relationships and ordering across cell populations, we performed a diffusion pseudotime (DPT) analysis, uncovering a hierarchical organization starting from actively proliferating RGCs and later segregating into 3 main branches (i.e. IN, EN1 and EN2) (Fig. 2b-d).

**Figure 2.**
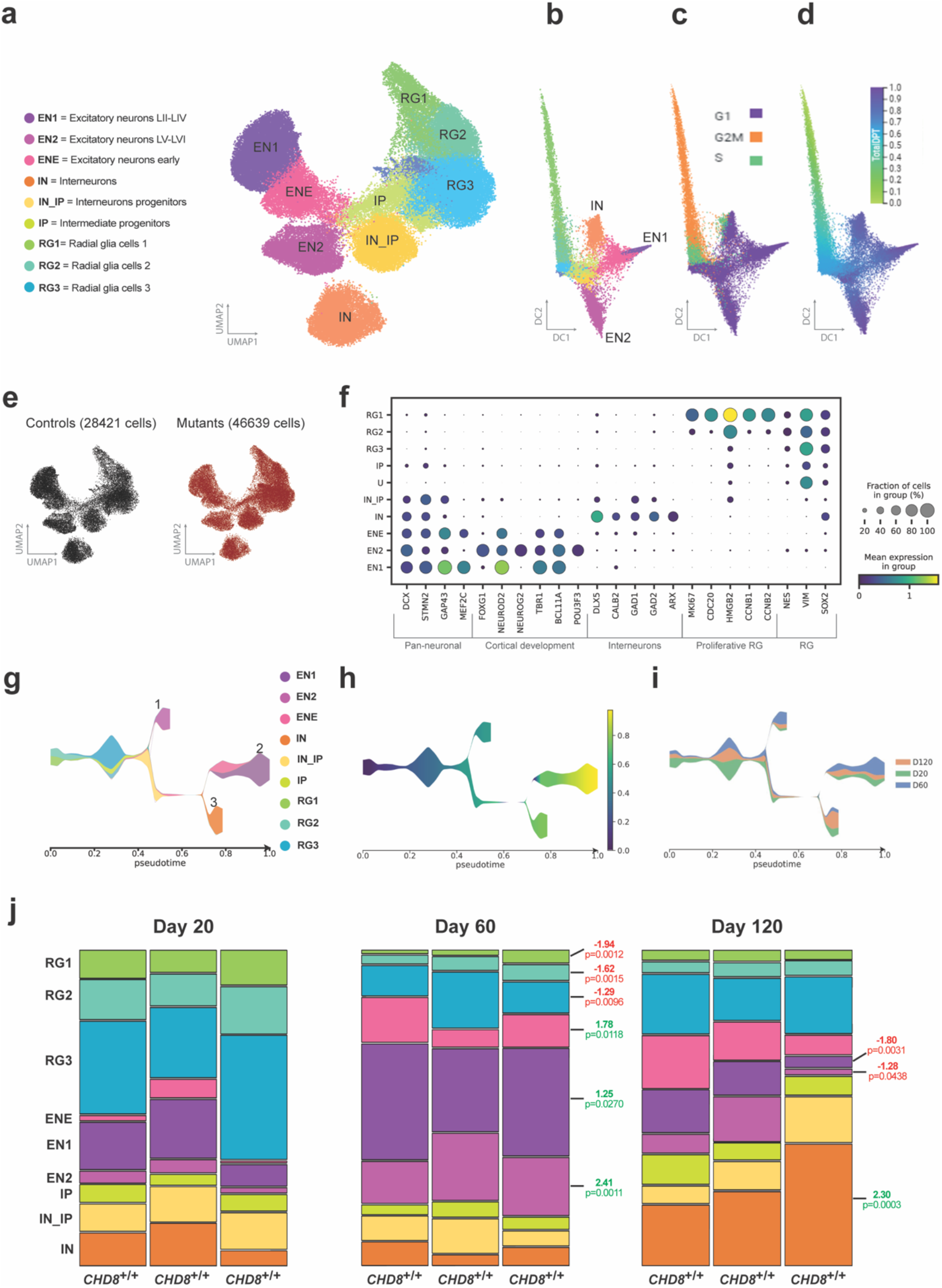
Cerebral organoids reproduce neurogenic lineages. (**a**) UMAP projection of droplet-based scRNAseq from cerebral organoids profiled at day 20, day 60 and day 120 *in vitro.* Coloured clusters show identified cell types. (EN1 = Excitatory neurons Layers II-IV, EN2 = Excitatory neurons Layers V-VI, ENE = Excitatory neurons early, IN = Interneurons, IN_IP = Interneuron intermediate progenitors, IP = Intermediate progenitors, RG1 = Radial glia cells 1, RG2 = Radial glia cells 2, RG3 = Radial glia cells 3). (**b**) Diffusion map depicting hierarchical organization of cell populations highlighting: (**c**) cell cycle stages and (**d**) pseudotime trajectories. (**e**) Total number of cells profiled per condition, 3 control (CTL) and 4 mutant (MUT) cell lines for each time point. Each sample was obtained by pooling 3 cerebral organoids obtained from at least two independent differentiation batches. (**f**) Expression enrichment of cortical neuronal classes canonical markers across clusters. (**g-i**) Tree graph, calculated in control lines, indicating distribution of cell populations into three main lineages connecting cycling radial glia with endpoints: 1) Excitatory neurons LIV-VI, 2) Excitatory neurons LII-III and 3) Interneurons. (**g**) Colour-code is according to cluster assignment, (**h**) total pseudotime calculated on all lineages or (**i**) time points. (**j**) Proportion of cells in each cluster across 3 controls lines divided by stage. P-Values and fold-change (red: down-regulated; green: up-regulated) are reported for populations identified as significantly changed in differential abundance analysis (see methods) comparing day 60 vs day 20 or day 120 vs day 60.

Analysis of population reproducibility across control organoids (obtained from independently grown clones of the original hESC line) corroborated the presence of comparable proportions of all identified subpopulations (Extended Data Fig. 5a,b) each of which followed the same reproducible pseudotemporal trajectory starting from early proliferating progenitors and bifurcating into inhibitory and excitatory lineages (Fig. 2b-d and Extended Data Fig. 5c), confirming that both population composition and developmental trajectories are robustly recapitulated across control organoid lines.

In order to corroborate with a complementary approach the reconstruction of the main developmental trajectories in control cerebral organoids (3 lines, n = 28421 cells), we applied tree graph analysis^30^ which recapitulated the three main developmental trajectories (EN1, EN2, IN). Each maturation trajectory is driven by the expression of specific transcription factors: 1) excitatory neurons Foxg1-Ngn2, 2) excitatory neurons NeuroD2, Tbr1 and 3) interneurons Dlx1/2 (Fig. 2g, Extended Data Figure 4b). These lineages displayed a stereotypical pseudotime progression from progenitors to neuronal populations (Fig. 2b, g, h) with each neuronal population matching the expected increase in abundance throughout stages (Fig 2i-j).

In order to investigate the impact of *CHD8* haploinsufficiency on cortical development, we focused on the three *CHD8* mutant lines whose phenotypes consistently resulted in macrocephaly-matching organoid overgrowth (*CHD8^+/−^*, *CHD8^+/−;GFP^* and *CHD8^+/E1114X^*). Analysis of relative population densities across conditions revealed stage-specific alterations caused by *CHD8* mutations in the abundance of different populations, including a delayed production of EN1-EN2 together with a robust anticipated generation of IN and IN_IP (Fig 3a). Indeed, following the developmental branches through pseudotime, we confirmed a higher representation of mutant cells in the interneuron branch at day 60 and in the excitatory at day 120 (Fig 3b). Differential abundance analysis of cluster frequencies confirmed the statistical significance of these differences and indicated that the increase at day 60 in the number of interneurons of the parvalbumin lineage (Fig 3c, Extended Data Fig. 6a,b) is the most robust population change caused by *CHD8* haploinsufficiency.

**Figure 3.**
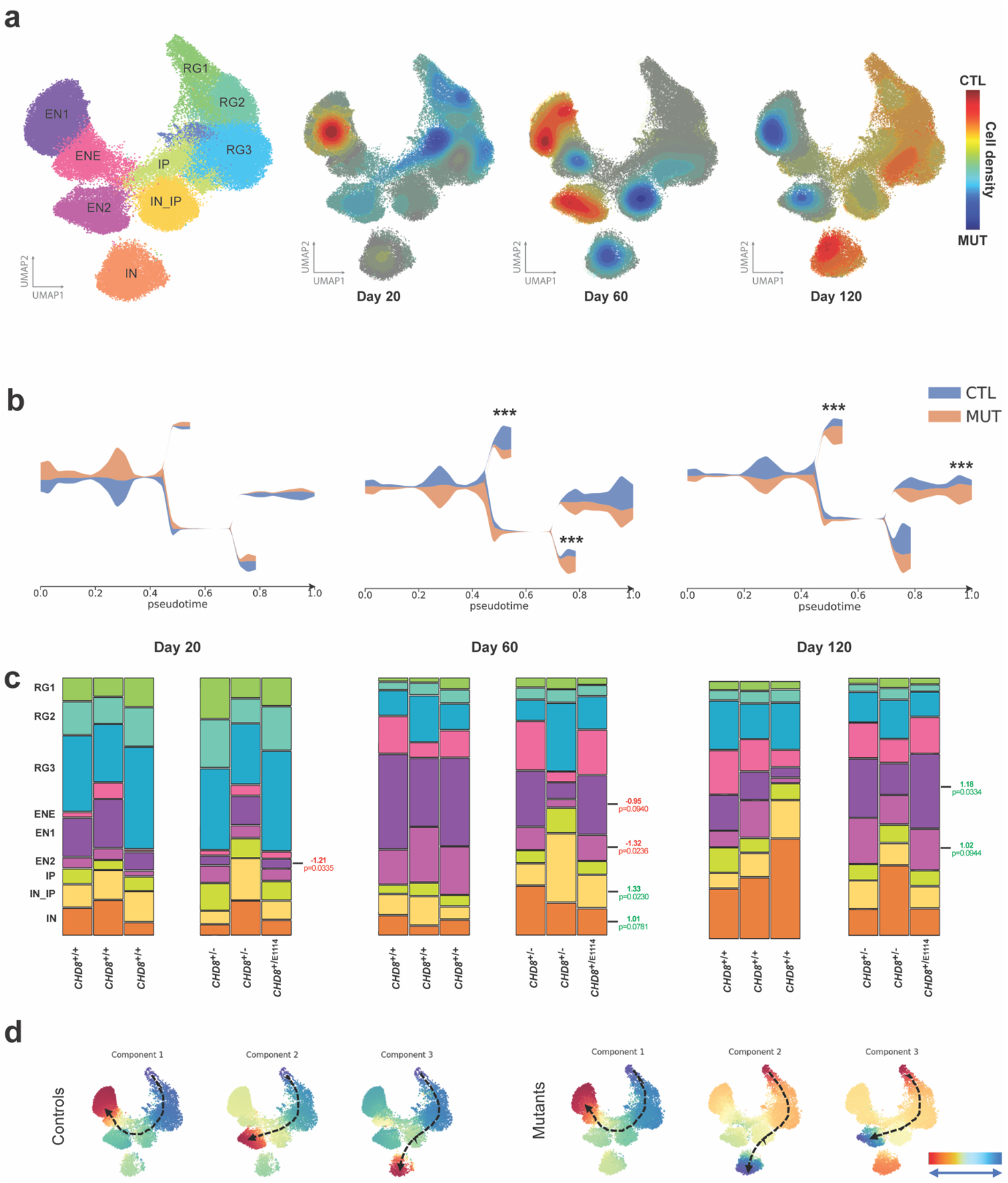
*CHD8* haploinsufficiency causes transient aberrations in cell population proportions and early interneuron differentiation. (**a**) Density counter plot highlighting stage-wise differences in cell frequencies (blue, high in mutant, red, high in control). It indicates specific sub-areas of cell clusters enriched in mutant or control conditions. (EN1 = Excitatory neurons LII-LIV, EN2 = Excitatory neurons LV-LVI, ENE = Excitatory neurons early, IN = Interneurons, IN_IP = Interneurons progenitors, IP = Intermediate progenitors, RG1 = Radial glia cells 1, RG2 = Radial glia cells 2, RG3 = Radial glia cells 3). (**b**) Tree graph displaying stage-wise differences in proportion of cells between control and mutant per branch (lineage). Asterisks indicate areas where frequency differences were significant, calculated as reported in panel **c**. (**c**) Proportion of cells in each cluster across conditions divided by stage. P-values and fold-changes (green: up-regulated; red: down-regulated) are calculated by differential abundance analysis (see methods) comparing mutant vs control organoids at each stage. (**d**) First three pseudotime components from control and mutant cerebral organoids at day 60, indicating the first three main trajectories within the populations, defined between the two ends (blue and red, arrows), highlighting a preference for interneuron differentiation in mutant organoids.

In order to understand whether this difference in population frequencies was a result of aberrant developmental trajectories, we calculated, separately for mutants and controls, the first three pseudotime components at each stage, thus revealing the three most likely trajectories in each condition at each stage. This analysis revealed that specifically at day 60, while in control organoids the first two most likely trajectories result in two different excitatory neuronal clusters, in *CHD8* mutant organoids the second most likely trajectory anticipates interneuron production (Fig. 3d), thus further indicating that *CHD8* haploinsufficiency favors accelerated interneuron development.

### *CHD8* haploinsufficiency causes sustained proliferation in human brain organoids

Next, we employed immunohistochemistry to both validate and dissect the mechanism of the delayed generation of excitatory neurons. To this end we established an unbiased method to quantify the amount of different cell types in cerebral organoids *in situ* by adapting a deep learning-based approach for semi-automatic counting of immunolabeled cells. In short, we performed automated cell body segmentation in manually curated regions of interest using Cellpose^31^, a state-of-the-art pre-trained deep neural network for cell segmentation that enabled us to analyze large portions of the organoids instead of focusing on small manual counting windows (Extended Data Fig. 7).

To evaluate if the increased size and delayed neurogenesis are due to a proliferative imbalance of neural progenitor cells, we pulsed *CHD8^+/+^* and *CHD8^+/−^* cerebral organoids at d10 and d20 with EdU for one hour and checked its incorporation into newly synthesized DNA after 16 hours. Interestingly, *CHD8^+/−^* organoids, at both developmental stages, displayed significantly more EdU and Ki67 positive cells than in the *CHD8^+/+^* samples (Fig. 4 a, b) indicating that already at day 10, and consistently at day 20, *CHD8* mutations are associated with an increased number of proliferating cells. This difference was not due to changes in cell cycle length as suggested by an equal ratio of Edu/Ki67 positive cells in mutant and control organoids 16 hours after the EdU pulse (Fig. 4 a, b). Interestingly, at day 10 *CHD8^+/S62X^* organoids do not show any difference in EdU positive cell number compared to control organoids (Extended Data Fig. 8), a finding maintained through day 20 (with only a non-significant trend towards increased EdU labelled cells). This observation is in line with the much milder phenotype observed in S62X mutant organoids and patients, indicating that the early increase in proliferating cells drives the observed increase in organoid size and macrocephaly in patients.

**Figure 4.**
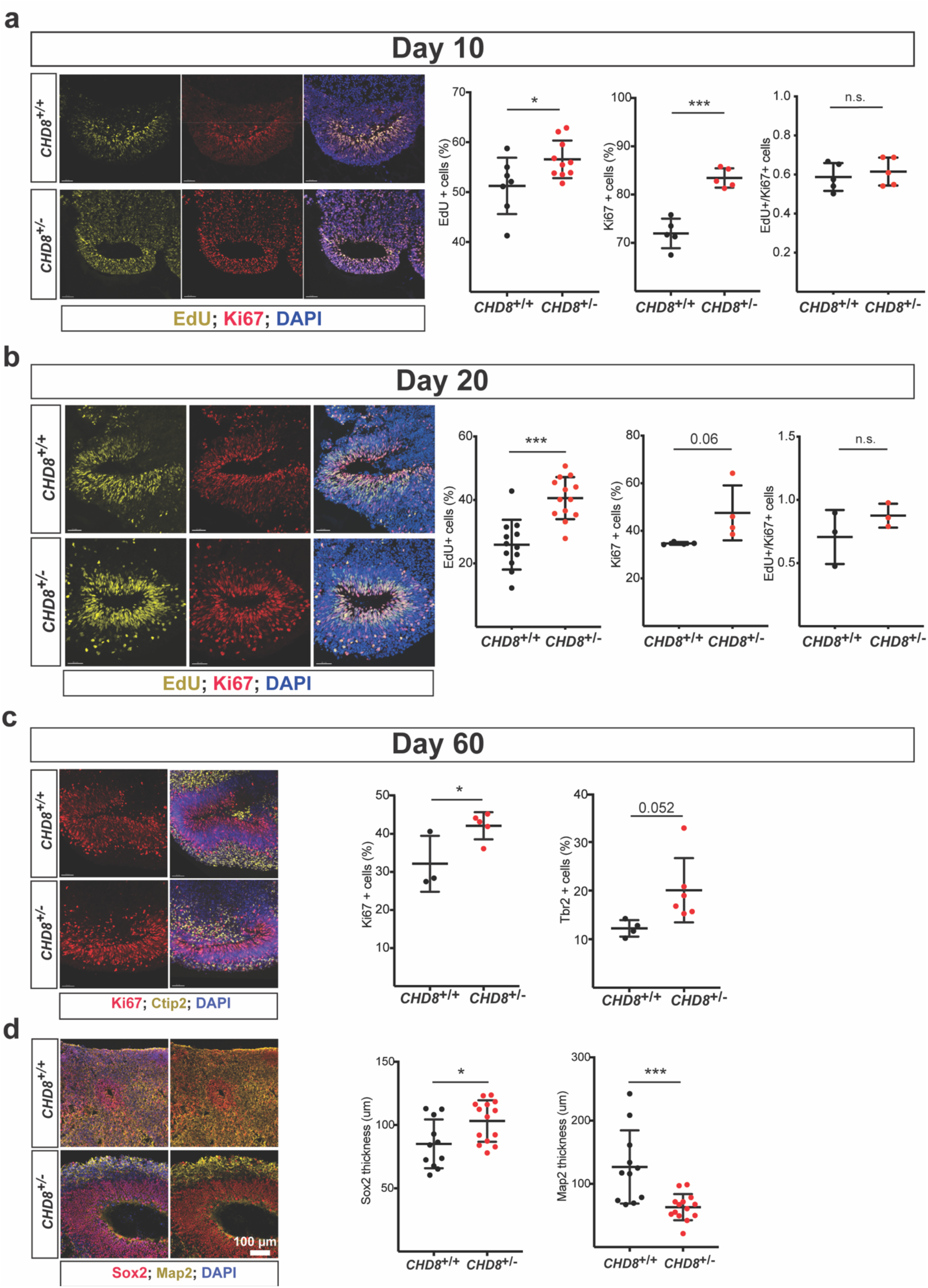
Increased and sustained proliferation in cerebral organoid carrying *CHD8* mutations. (**a**) Representative images of ventricular zone (VZ)-like structures of 10-day-old, *CHD8^+/+^* and *CHD8^+/−^*, cerebral organoids. Organoids were pulsed with BrdU for one hour and fixed 16 hours later. Sections were stained for BrdU (yellow), Ki67 (red) and DAPI (blue). Quantification (on the right) revealed a higher number of BrdU and Ki67 positive cells, but no difference in their ratio, in *CHD8^+/−^* organoids. The number of BrdU or Ki67 positive cells is represented as percentage of DAPI stained cells. (Unpaired t-test, *P < 0.01; ***P< 0.001; *CHD8^+/+^*, *n=7; CHD8^+/−^*, *n= 10 for BrdU; CHD8^+/+^*, *n=5; CHD8^+/−^*, *n= 5 for Ki67*). (**b**) Representative images and quantification of EdU (yellow) and Ki67 (red) double labelling of 20-day-old *CHD8^+/+^* and *CHD8^+/−^* organoids. Quantification confirmed an increased proportion of EdU and Ki67 positive cells in *CHD8^+/−^* VZ-structures (Unpaired t-test, ***P< 0.001 for EdU and P=0.06 for Ki67*; CHD8^+/+^*, *n=12; CHD8^+/−^*, *n= 13 for EdU; CHD8^+/+^*, *n=4; CHD8^+/−^*, *n= 4 for Ki67*). (**c**) Representative images of 60-day-old organoids stained for Ki67 (red) and Ctip2 neurons (yellow). Quantification (right panels) revealed an increased number of Ki67+ and Tbr2+ (images not shown) cells in *CHD8* mutant organoids (Unpaired t-test, *CHD8^+/+^*, *n= 3*; *CHD8^+/−^*, *n= 5 for Ki67; CHD8^+/+^*, *n= 4*; *CHD8^+/−^*, *n= 6 for Tbr2*). (**d**) Representative images for cortical structures from day 60 *CHD8^+/+^* and *CHD8^+/−^* cerebral organoids stained for Sox2 (red) and Map2 (yellow). Quantification (right panel) revealed a significant increase in Sox2+ layer thickness and a significant decrease in Map2+ layer thickness (Unpaired t-test, *CHD8^+/+^*, *n= 11*; *CHD8^+/−^*, *n= 13*). *P < 0.05; **P < 0.01; ***P < 0.001; ****P < 0.0001; n.s., non-significant. n, number of organoids investigated (multiple rosettes per organoids were analysed) obtained from at least 2 independent batches. Results are presented as mean ± s.d‥

In line with the expansion of the proliferative compartment, we found that at the same early stages the number of Tbr2-positive intermediate progenitors is decreased in the *CHD8^+/−^* organoids (Extended Data Fig. 9). This reduced neuronal output was reflected at later time points in decreased thickness of the neuronal layer, as identified by Map2 staining, and increased amounts of Sox2 positive radial glia progenitor cells. Together, our data argue that macrocephaly-associated *CHD8* haploinsufficient mutations lead to increased brain size in humans by extending the phase of neural stem cell proliferation, increasing the progenitor pool and eventually resulting in the generation of a higher number of neurons at later stages.

### *CHD8* haploinsufficiency affects neural stem cell proliferation cell-autonomously

To understand whether *CHD8* haploinsufficiency effects are cell-autonomous while controlling for any possible confounder due to the inherent variability of the model, we employed the stable eGFP-expressing *CHD8* LoF mutant hESC line (*CHD8^+/−;GFP^*) (Fig. 1d) to generate mosaic organoids allowing us to track mutant cells at all stages of maturation. We generated control (*CHD8^+/+^*), GFP-*CHD8* mutant (*CHD8^+/−;GFP^*) and mixed (1:1) control/GFP-mutant (*CHD8^+/+^*/*CHD8^+/−;GFP^* hereinafter referred to as *CHD8*^mix^) organoids (Fig. 5a). We next counted the proportion of eGFP positive or negative progenitor (Sox2 positive) and neuronal (NeuN positive) cells at day 60 in control, mutant and mixed organoids. In line with the thicker Sox2 layer observed in the *CHD8^+/−^*organoids (Fig. 4d) and indicating a cell autonomous phenotype in mixed organoids at day 60, we found a higher proportion of Sox2+/GFP+ cells (i.e. *CHD8^+/−^*), compared with Sox2+/GFP-cells (i.e. *CHD8^+/+^*) (Fig. 5b). Conversely, at the same stage we observed a lower proportion of *CHD8* mutant neuronal cells (i.e. NeuN+/GFP+) compared to wild type cells (i.e. NeuN+/GFP-) (Fig. 5b).

**Figure 5.**
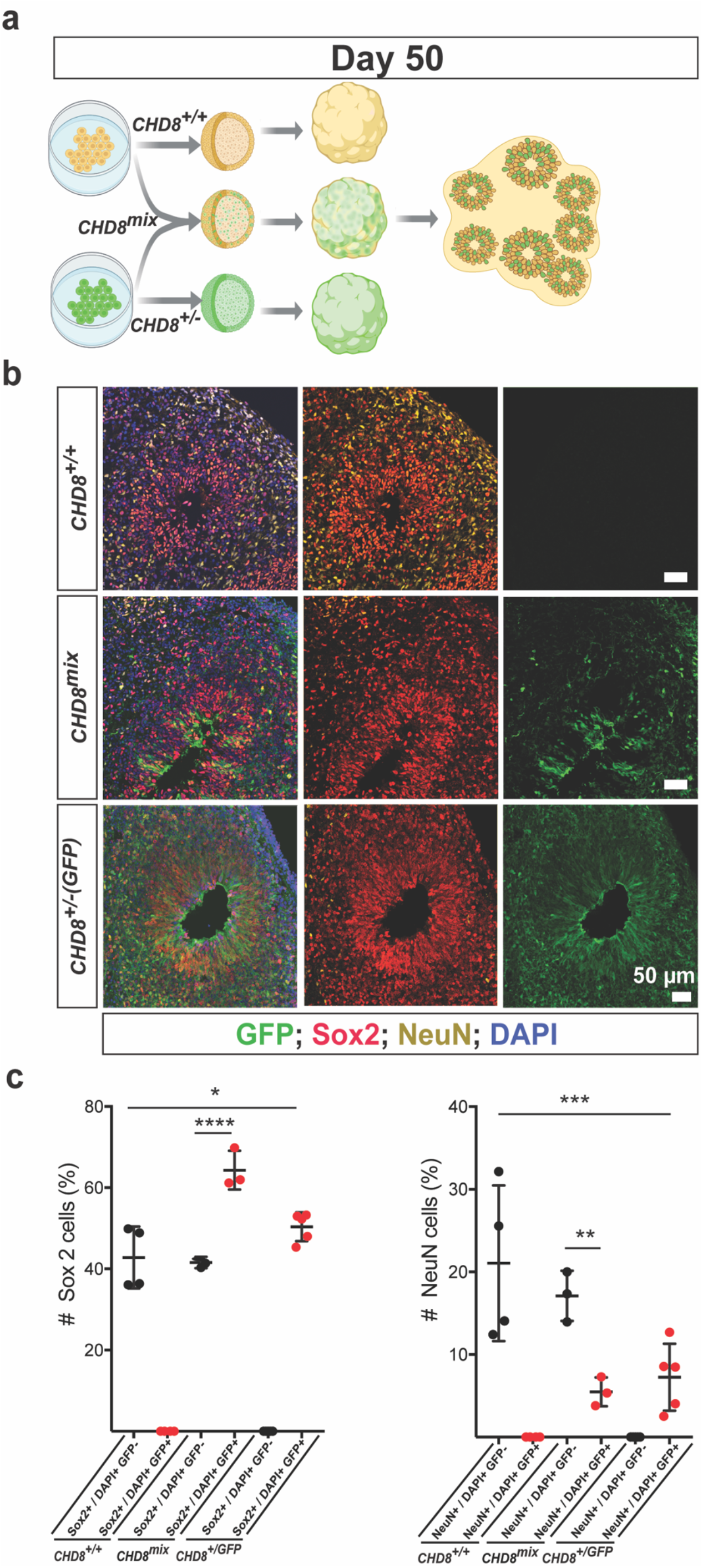
*CHD8* haploinsufficiency leads to cell-autonomous defects. (**a**) Cartoon explaining the strategy for the generation of *CHD8^+/+^*, *CHD8^+/+^* 1:1 mixed with *CHD8^+/GFP^* (*CHD8^mix^*) and *CHD8^+/GFP^* cerebral organoids. (**b**) Representative images of *CHD8^+/+^* and *CHD8^mix^* and *CHD8^+/−;GFP^* developing cortical structures, stained for the neuronal marker NeuN (yellow) and Sox2 at day 60. (**c**) Quantifications revealed a significant increase of Sox2+ cells in *CHD8^+/GFP^* and a larger proportion of Sox2+GFP+ cell population in *CHD8^mix^* cortical structures. (Ordinary One-way ANOVA followed by Dunnett’s multiple comparisons test; *CHD8^+/+^*, *n= 4; CHD8^+GFP^*; *n=3; CHD8^mix^, n=5*). Furthermore, a significant reduction in NeuN+ neurons among *CHD8^+/GFP^* cells was observed in the *CHD8^mix^* and in *CHD8^+/GFP^* cortical structures. (Ordinary One-way ANOVA followed by Dunnett’s multiple comparisons test; *CHD8^+/+^*, *n= 4; CHD8^+/GFP^*, *n=3; CHD8^mix^, n= 5*). *P < 0.05; **P < 0.01; ***P < 0.001; ****P < 0.0001. n, number of organoids investigated (multiple rosettes per organoids were analysed). Results are presented as mean ± s.d‥

Together, these data indicate that heterozygous *CHD8* LoF mutations lead to a cell-autonomous sustained proliferation of neural precursors in human cerebral organoids.

### *CHD8* mutations alter proliferative gene expression program

Given the molecular function of *CHD8^32^* and the cellular phenotypes observed in *CHD8* mutant organoids, we carried out a transcriptomic analysis on 10 day old cerebral organoids, a stage enriched for proliferating cells that marks the first deviations between *CHD8^+/+^* and *CHD8^+/−^* cerebral organoid development (Fig. 4 a). Differential expression analysis revealed 868 genes dysregulated as a result of *CHD8* LoF, comprising in equal amounts downregulated and upregulated genes (Extended Data Table 3). Consistent with previous reports hypothesizing functional convergence of ASD-associated genes around similar stages of human neocortical development^2,3^, as well as with experimental data revealing important co-expression networks around *CHD8* targets in human midfetal brain and neural stem cells, we observed a significant enrichment in top ASD-associated genes among the differentially expressed genes (DEGs) (Fig. 6a and Extended Data Table 3), including other high-risk genes associated with chromatin regulation such as *ASH1L* and two established direct targets of CHD8, *ARID1B* and the NCoR-complex subunit *TBL1XR1*. Gene Ontology (GO) analysis showed upregulated genes to be enriched for terms related to cell cycle regulation, more specifically for genes implicated in G1/S phase transition, RNA splicing and regulation of transcription (Fig. 6b, Extended Data Table 3). In stark contrast, downregulated genes were enriched for terms related to neuronal differentiation and brain development (Fig. 6b, Extended Data Table 3), underscoring how already at early stages of human cortical development *CHD8* haploinsufficiency alters the regulation of neural progenitor proliferation and neuronal differentiation. Furthermore we found that, among the down- and up-regulated genes, respectively 28% and 47% are reported as CHD8-bound^33^ (Fig. 6c), pointing to a significant direct effect of *CHD8* haploinsufficiency. Interestingly, GO term enrichment analysis on downregulated genes showed a pronounced functional partitioning between direct and indirect targets, with indirect targets enriched in neurogenesis regulation while direct targets were instead enriched for chromatin modifiers and transcriptional regulators (Fig. 6d, Extended Data Table 3). This suggests that CHD8 acts directly on gene regulation in progenitor cells while the observed downregulation of neurogenesis pathways is indirect, reflecting either intermediate effectors or the proliferative skew of the neural stem cell population. Upregulated genes on the other hand showed no marked difference in GO term enrichment between direct and indirect targets (Extended Data Table 3) but rather recapitulated the enrichment of cell cycle terms.

**Figure 6.**
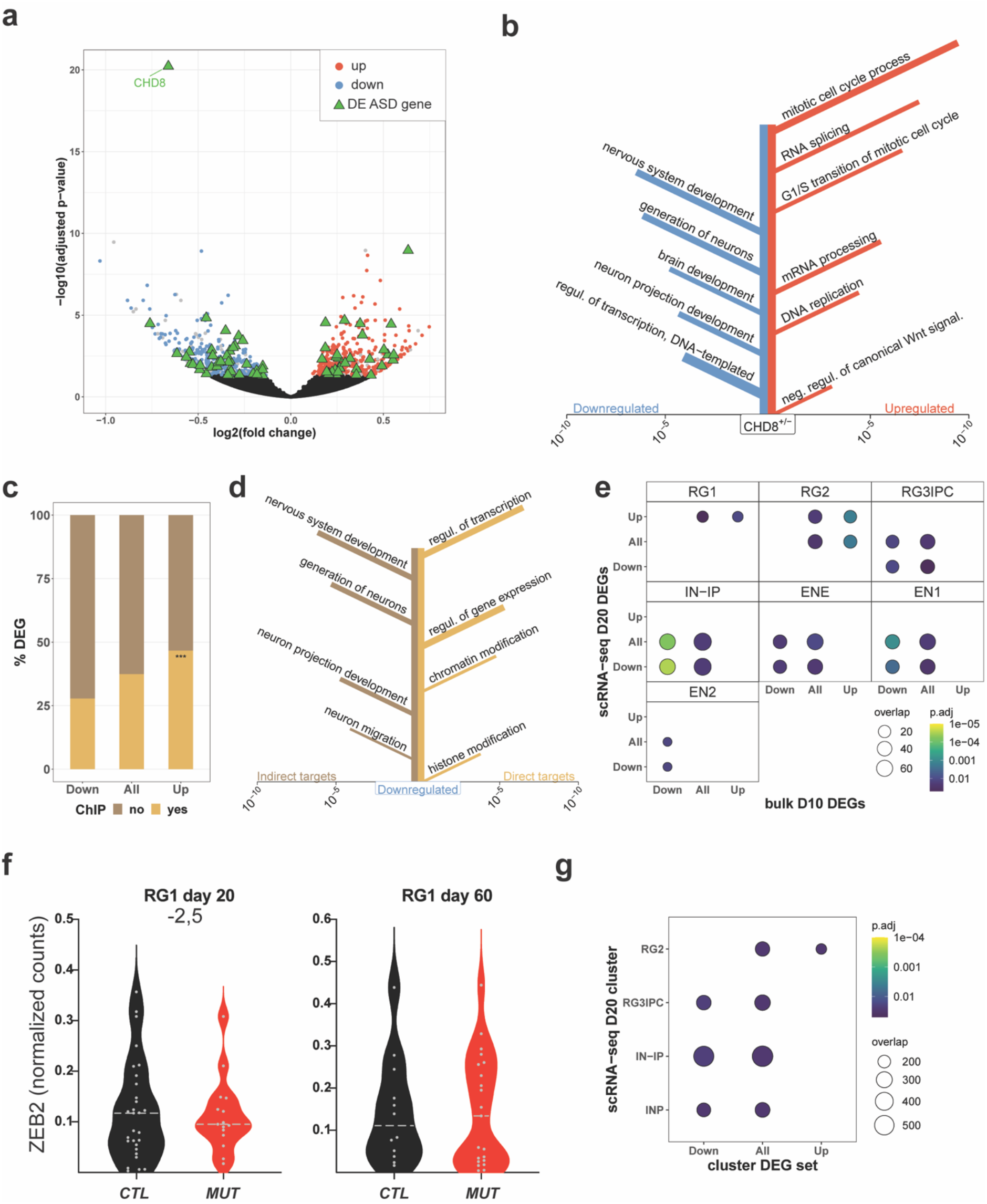
Cell-type specific gene expression defects in *CHD8* mutant cerebral organoids. (**a**) Bulk RNA sequencing of day 10 organoids reveals overall moderate fold changes in gene expression upon haploinsufficiency of *CHD8*. The volcano plot displays the results from the differential expression analysis between *CHD8^+/+^* and *CHD8^+/−^* organoids as negative decadian logarithm of the FDR adjusted p-value on the y-axis and the binary logarithm of the fold change on the x-axis. Genes displaying significant differential expression between *CHD8^+/+^* and *CHD8^+/−^* (adjusted p-value <=0.05, 868 genes) are shown in blue and red for genes down- and upregulated in *CHD8^+/−^* organoids, respectively. Genes excluded because of significant differences between the two wildtype controls are shown in grey and genes associated with ASD based on the SFARI list are displayed as green triangles (Extended Data Table 3). (**b**) The graph shows selected Gene Ontology (GO) biological processes enriched in genes downregulated (left) or upregulated (right) in *CHD8^+/−^* organoids. Branch-length represents significance of the enrichment while branch-thickness displays the number of dysregulated genes associated with each term. (**c**) Overlap between our list of 868 dysregulated genes and CHD8 targets based on published ChIP-seq data^33^ shows significant enrichment for CHD8 binding in upregulated genes (***P < 0.001, Fisher’s exact test, Extended Data Table 3). (**d**) GO term enrichment for downregulated genes split into direct and indirect targets. (**e**) Overlap between DEGs identified in bulk RNA-seq from day 10 and DEGs from scRNA-seq at day 20 (no FC cutoff). Comparisons are for all DEGs as well as down- and upregulated DEGs separately. Only significant overlaps (Fisher’s exact test FDR adj. p-value < 0.05) are shown. (**f**) ZEB2 expression in RG1 at day 20 and day 60 from scRNA-seq data. At day 20, but not at day 60, ZEB2 is significantly downregulated in mutant samples. (**g**) Overlap between ZEB2 target genes (see methods) and DEGs in scRNA-seq at day 20 (no FC cutoff). Different clusters are shown on the y-axis, the x-axis displays which set of DEGs was used for the overlap (all, upregulated, or downregulated). Only significant overlaps are included (Fisher’s exact test FDR adj. p-value < 0.05).

We then extended the analysis of CHD8 dosage-dependent transcriptional dysregulation at the single cell level by stage-specific differential gene expression analysis comparing macrocephaly-associated mutants versus control lines and thereby cross-validating bulk and single cell RNAseq. We found significant overlap between the DEGs identified through the two transcriptomic readouts and, importantly, the dichotomy between proliferation and neurogenesis enrichments observed in bulk data at day 10 was corroborated by the findings that upregulated and downregulated genes mapped onto DEGs in different day 20 cell clusters (Fig. 6e), with the former enriched in clusters of cycling radial glia cells while the latter were enriched in clusters further along the differentiation path.

We thus proceeded to annotate the function of genes aberrantly expressed in *CHD8* mutants in specific cell types, performing GO enrichment analysis for each cluster at each developmental stage (Extended Data Table 2). This analysis revealed an upregulation of genes belonging to GO terms related to cell cycle, mRNA metabolism, ribosome biogenesis and translation initiation (Extended Data Table 2) in proliferating cells (RG1-3). Next, to identify candidate regulators for the prolonged proliferative phase in *CHD8* mutants, we focused on the most highly significant DEGs specifically retrieved from the clusters of day 20 proliferating cells and found ZEB2 to be severely down-regulated in *CHD8*-mutant RG1 cells (2,5-fold below control levels, Fig. 6 f). ZEB2 encodes a transcription factor recently described to promote neuroepithelial differentiation into radial glia, thus determining the initial number of proliferating cells and thereby controlling cortical tissue expansion. For this reason, the delayed increase in ZEB2 levels in humans versus apes has been proposed to underlie human neocortical expansion^34^. Consistently, we found enrichment for ZEB2 ChIPseq-defined targets among the DEGs of multiple clusters at day 20 (Fig. 6g), whose specificity (enrichment in upregulated genes for cycling cells and for downregulated genes in more differentiated cells) further implicates ZEB2 as a critical mediator of the impact of *CHD8* haploinsufficiency on the proliferative dynamics of the neural stem cell compartment.

### *CHD8* mutations affect mRNA metabolism in post-mitotic neurons

Finally, when plotting the number of DEGs per cluster per developmental stage we noticed that both early corticofugal neurons (ENE and IN_IP) and inhibitory neuron progenitors show the highest number of DEGs (Figure 7a), while some cell types show nearly no DEGs at any of the three chosen time points (Figure 7a). We thus focused on those clusters with the highest number of DEGs to elucidate the downstream impact of *CHD8* haploinsufficiency on human cortex developmental programs, noting that two of the most affected clusters in terms of DEGs (ENE and IN_IP) are the ones immediately preceding the most affected clusters in terms of cell frequencies (EN1 and IN) (Fig. 3), thereby linking upstream transcriptional changes to downstream population imbalances. In light of previous findings from murine fetal brain that associated alternative splicing alterations to CHD8-associated developmental delay^14^, we investigated whether RNA processing could also be impacted in the human setting and related to specific developing subpopulations. We thus probed the delta percent spliced-in index (dPSI)^35^ of expressed genes in day 20 IN_IP, IN and ENE/EN1 as a readout for changes in splicing patterns. As a result, we found significantly different dPSI scores in a subset of genes in IN_IP, IN and ENE/EN1 populations when comparing CHD8^+/+^ and CHD8^+/−^ and benchmarking against the physiological regulation of alternative splicing across control clusters. Strikingly, functional enrichment analysis of these subsets pointed to mRNA metabolic processing in the IN_IP and IN populations (the latter also returning a specific enrichment for splicing) while no significant functional enrichment was detected in the ENE/EN1 population (Figure 7b). These findings uncover a dysregulation cascade that amplifies, selectively in the interneuron branch, the primary *CHD8* dosage-dependent effects on alternative splicing into further RNA processing alterations.

**Figure 7.**
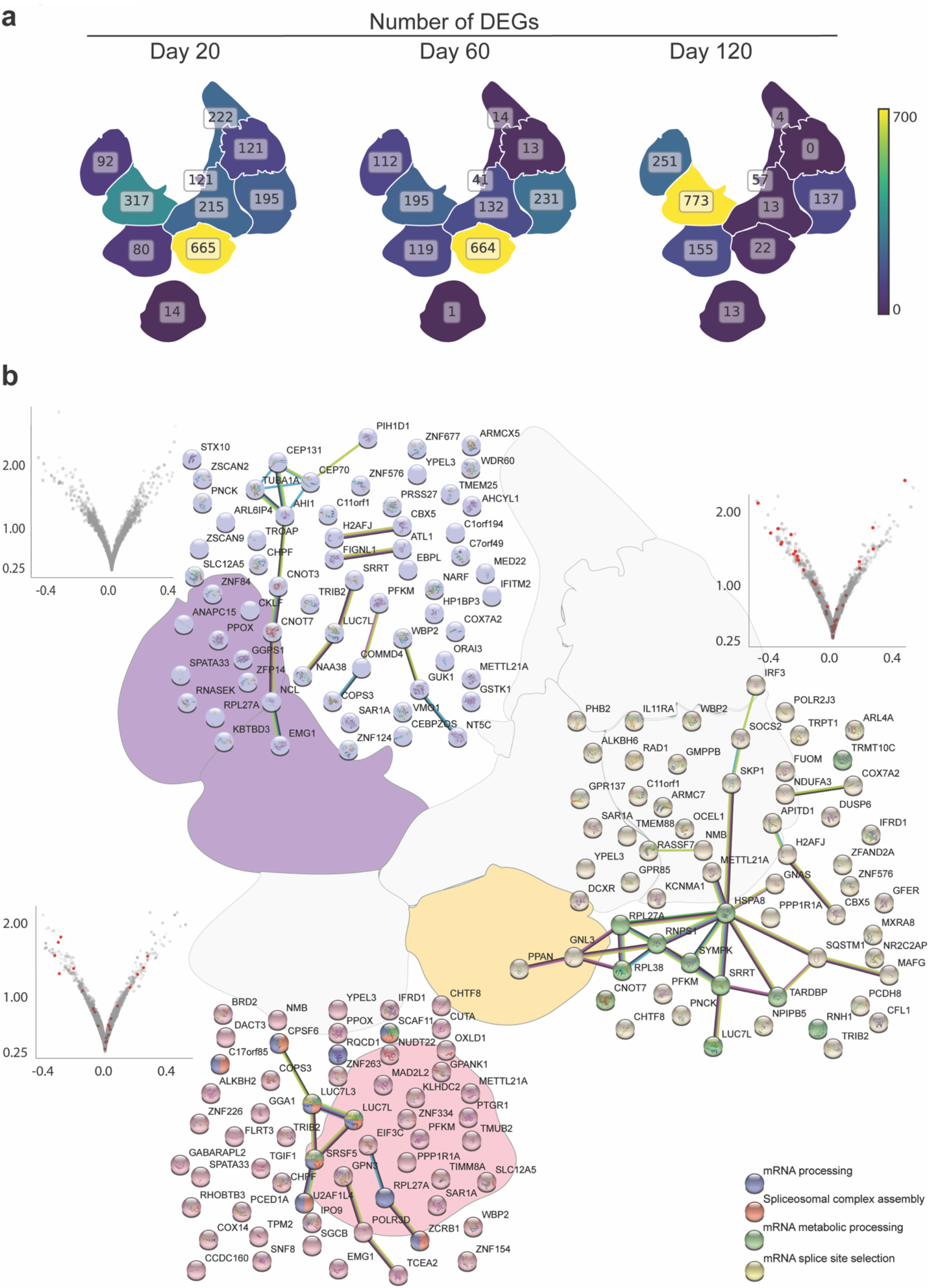
*CHD8* haploinsufficiency leads to changes in the transcriptional landscape of specific cell types. **(a)**UMAP outline illustrating for each cluster and stage the number of DEGS (FDR<0.05 and log2FC>|1|) resulting from mutant versus control comparison. **(b)** Single cell RNA sequencing points to differences in alternative splicing in *CHD8* mutants with changes at the level of transcripts of a given gene present in ENE/EN1, IN and IN_IP clusters at day 20. For these clusters, the volcano plot shows the results of the differential splicing analysis by displaying the negative logarithm of the adjusted p-value on the y-axis and the binary logarithm of the Delta Percent-Spliced-In (dPSI) on the x-axis. Genes displaying significant differences in splice site usage between *CHD8^+/+^* and *CHD8^+/−^* (p-value <=0.05, ENE/EN1: 67 genes, IN_IP: 65 genes and IN: 62 genes) are depicted as an interaction network, where the nodes represent proteins and the edges evidence of functional or physical interactions from different sources indicated by the edge colour-code: known interactions in cyan (curated databases) or magenta (experimental evidence); predicted interactions in red (gene fusion), green (gene neighborhood), blue (gene co-occurrence), yellow (text-mining), black (co-expression) and lilac (protein homology). Genes with significant differences between wildtype controls were excluded from the analysis. Nodes are coloured according to gene ontology categories resulting from Gene Ontology (GO) functional enrichment analysis within the network (biological process domain) that reached significance (FDR, adjusted p-value <=0.05, Benjamini– Hochberg correction). IN_IP: GO:0016071 – RNA metabolic process (red), adj. p-value=0.01; IN:GO:0000245 – spliceosomal complex assembly (red), adj. p-value= 0.03, GO:0006397 – mRNA processing (blue), adj. p-value= 0.03, GO:0016071 – mRNA metabolic processing (green), adj. p-value=0.03, GO:0006376 – mRNA splice site selection (yellow), adj. p-value=0.04. The same genes are highlighted as red dots in the Volcano Plot.

## Discussion

As one of the most prevalent and penetrant causes of ASD, *CHD8* haploinsufficiency is paradigmatic, for NDD, of the major gap that exists between the increasing resolution of genotype-phenotype relationships and the virtually complete lack of knowledge of the human neurodevelopmental trajectories that underlie them. Thus, while ASD features are invariably present, albeit with variable severity, macrocephaly is present in only up to one half to two thirds of cases^36^, affording the opportunity to dissect the impact of different mutations in terms of shared versus distinct neurodevelopmental outcomes. Here we pursued a first genotype-phenotype dissection of *CHD8* haploinsufficiency in terms of human neurodevelopmental trajectories at single cell resolution.

First, we focused on the initial stages of neuroectoderm specification and modified culture conditions to improve both the efficiency and reproducibility of cerebral organoid derivation as a prerequisite for robust phenotypic scoring. We next built on our previous transcriptomic benchmarking of iPSC-based disease modelling^24^ and adopted a longitudinal isogenic design aimed at:i) comparing in the same genetic background LoF of the critical CHD8 helicase domain with patient-specific truncating mutations differentially associated to macrocephaly; ii) tracking CHD8 dosage-dependent neurodevelopmental trajectories by sampling multiple time-points corresponding to key morphological and molecular milestones; iii) resolving the cell-autonomous component of *CHD8* haploinsufficiency.

Our findings enable to draw the following conclusions. First, *CHD8* haploinsufficiency alters the proliferation/differentiation dynamics, with an early expansion of the neural progenitors’ compartment. Control-mutant mosaic organoids show this effect to be cell-autonomous, while both bulk and single-cell transcriptomics converged on the identification of its molecular underpinnings in terms of a pervasive dysregulation, in radial glia cells, of neuronal differentiation and cell cycle regulatory pathways. These were, respectively, down- and up-regulated and included a significant fraction of bona-fide CHD8 direct targets. Importantly, these early CHD8 dosage-dependent targets overlap significantly with bona fide ASD-causative genes^6^, pointing to a major layer of convergence and paving the way to future studies that can now systematically benchmark this convergent subset of ASD syndromes against the neurodevelopmental trajectories uncovered in this work. Importantly, while the relevance of the genetic background even for highly penetrant mutations is being increasingly recognized^37^, the observation that the patient-specific mutation not associated to macrocephaly did not induce overgrowth underscores the sensitivity of our organoid system in capturing pathophysiological readouts of patient-specific mutations.

Second, the *CHD8* mutations causing cerebral organoid overgrowth (and associated to macrocephaly in patients) disrupt the balance between excitatory and inhibitory neuronal production, with a delayed production of excitatory neurons (both lower and upper layers) and a major increase in interneuron output at day 60 that emerged from single cell transcriptomics as the most conspicuous subpopulation change in mutant lines. This temporal and lineage specificity of *CHD8* impact acquires even greater salience through the partitioning of gene expression dysregulation revealed by single cell transcriptomics. Indeed, only selected cell populations appear vulnerable to CHD8 dosage, and among these only some significantly so throughout development, as in the case of the RG3 cluster of radial glia (corresponding to radial glia in the G1 phase of the cell cycle) and the corticofugal neurons. The contrast is indeed particularly striking when comparing the excitatory and inhibitory trajectories, with interneuron progenitors and early intermediates greatly affected only through day 60, while both lower and upper layer excitatory neurons become increasingly affected over time. These findings uncover a bimodal developmental impact of *CHD8* haploinsufficiency, with an early switch that accelerates interneuron fate acquisition but with relatively little lasting impact on interneuron transcriptional identity, accompanied by a delay in excitatory fate that leaves a major lasting legacy on the transcriptional identity especially of lower layer neurons.

Finally, by probing cell type-specific transcriptional dysregulation, we uncovered RNA processing as a major layer of CHD8 dosage-dependent vulnerability. Mouse studies had previously reported splicing alterations in *Chd8* heterozygous mutants and related them to developmental delay^14^. Here we found that in mutants early interneuron and excitatory compartments as well as interneurons displayed a significant difference in the pattern of alternatively spliced transcripts. Strikingly, these alternatively processed transcripts specifically recovered from the cell clusters most impacted by *CHD8* haploinsufficiency code in turn for proteins involved in mRNA processing and splicing, pointing to an amplifying cascade of CHD8 dosage-dependent dysregulation in RNA processing. Importantly, having included in the analysis the physiological dynamics of developmentally regulated alternative splicing in controls, we can rule out that the aberrations in mutants are a byproduct of developmental skews rather than a primary effect of *CHD8* haploinsufficiency. In conclusion, our work exposes the neurodevelopmental impact of CHD8 haploinsufficiency, highlighting its exquisite specificity in terms of affected pathways and temporal windows that underlie patient-relevant endophenotypes.

## Acknowledgments

We thank Farnaz Freeman for technical assistance. This research was supported by the Scientific Service Units (SSU) of IST Austria through resources provided by the Bioimaging Facility (BIF) and the Life Science Facility (LSF). This work supported by the European Union’s Horizon 2020 research and innovation program (ERC) grant 715508 to G.N. (REVERSEAUTISM) and grant 825759 to G.T. (ENDpoiNTs); the Fondazione Cariplo 2017-0886 to A.L.T.; the Austrian Science Fund FWF I 4205-B to G.N‥

## Authors contributions

B.O., R.S., A.C.Y., J.M., generated and biochemically characterized cell lines and cerebral organoids, C.E.V. and C.C. performed the computational analysis of the single cell datasets. C.E.V., A.L.T., C.C., annotated single cells datasets, A.L.T. and Mi.G. performed single cell experimental procedures, C.P.D, performed and analyzed bulk transcriptomic experiments, C.S. developed semi-automatic cell counting, C.E.V., A.L.T., C.C., G.N. and G.T. developed the interpretive framework of the study. G.N. and G.T. led the study.

