## Supplementary Figures for "*CHD8* haploinsufficiency alters the developmental trajectories of human excitatory and inhibitory neurons linking autism phenotypes with transient cellular defects"

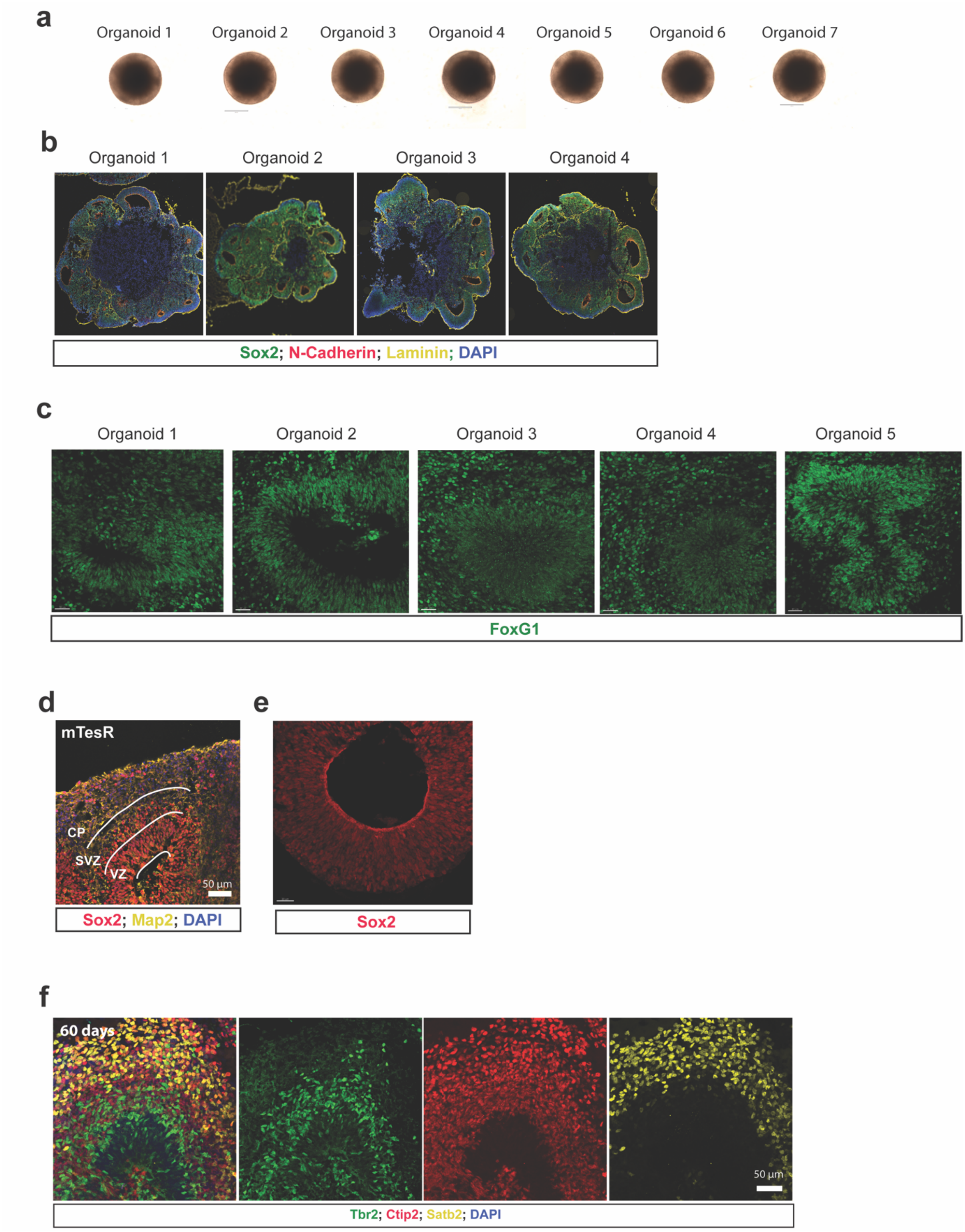


**Extended Data Figure 1. Characterization of cerebral organoids.** (**a**) representative pictures of cerebral organoids at day 0. Note the reproducible brightening of the neuroectoderm. (**b**) Representative images of cerebral organoids at day 10. (**c**) Expression of the forebrain marker FOXG1 in Day 60 cerebral organoids. FOXG1 positive cells were found in all the rosettes of all the organoids stained with this marker. (**d-f**) Representative images of developing cortical structures of cerebral organoids. VZ= ventricular zone-like structure, SVZ= subventricular zone-like structure, CP= cortical plate-like structure and expression of different marker, including Sox2 (progenitors), Map2 (neurons). Note that the layers of developing cortical regions of mature *CHD8*$^{+/+}$ cerebral organoids at day 60 (**f**) are composed of intermediate progenitors (Tbr2+, green), deep layer neurons (Ctip2+, red) and upper layer neurons (Satb2+, yellow).

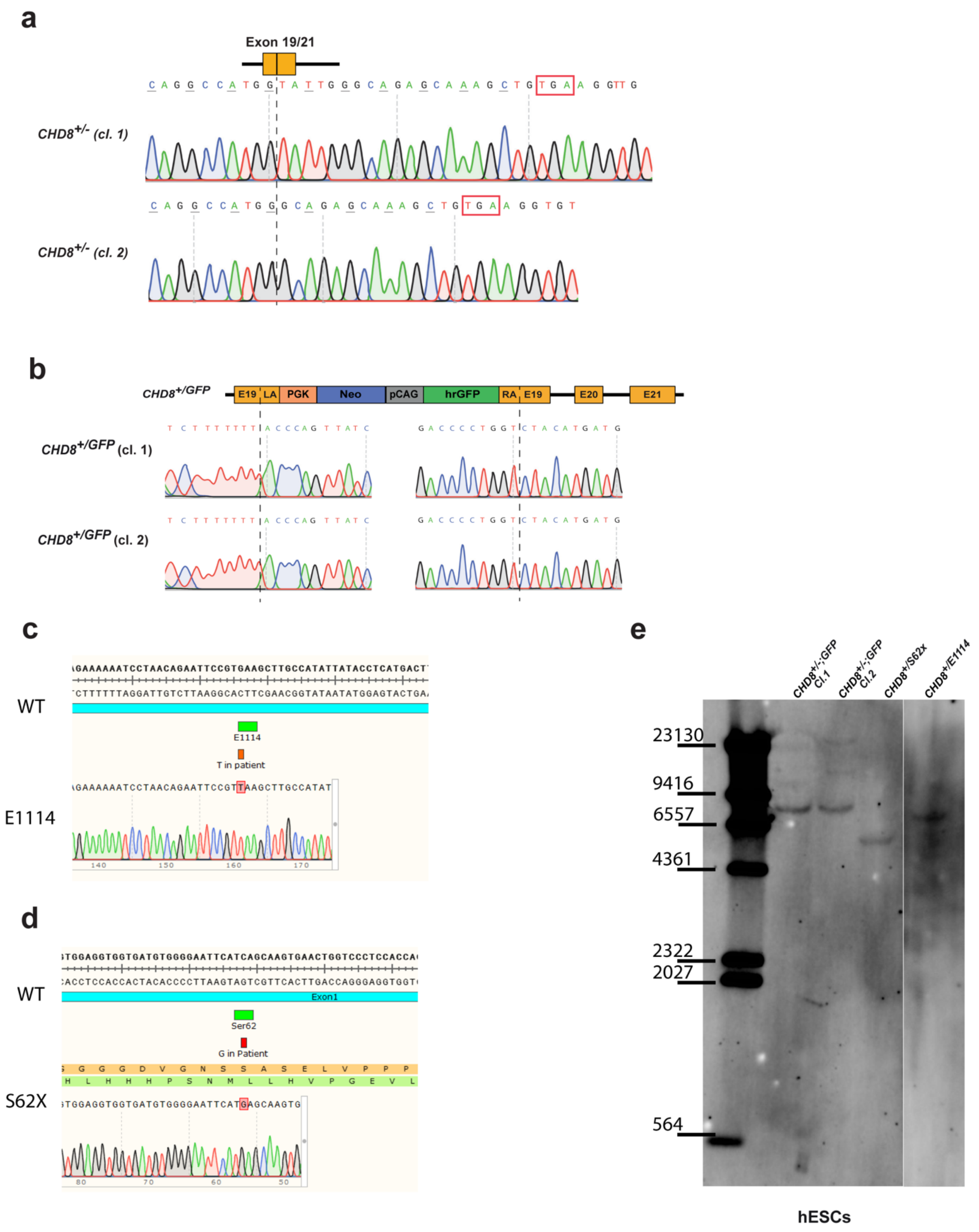


**Extended Data Figure 2. Generation of *CHD8* mutant hESC lines**. Schematics, southern blots and sequence chromatograms illustrating the CRISPR/Cas9-mediated generation of the *CHD8*$^{+/-}$ (**a**), *CHD8*$^{+/-;GFP}$ (**b**), *CHD8*$^{+/E1114;GFP}$(**c**), and *CHD8*$^{+/S62X;GFP}$ (**d**) hESC lines. (**a**) *CHD8*$^{+/-}$ cells were generated by deleting part of exon 19, whole exon 20 and part of exon 21, employing two guide RNAs (gRNA 1 and gRNA 2) targeting exon 19 and exon 21, respectively. For each mutation multiple clones were sequenced and analysed. Shown are sequence chromatograms spanning the site of exon 20 deletion in *CHD8*$^{+/-}$ clones (cl. 1 and cl. 2). (**b**) *CHD8*$^{+/-;GFP}$ clones were generated by CRISPR/Cas9- and HDR-mediated insertion of pCAG-hrGFP into exon 19 of *CHD8*. Shown are the sequence chromatograms for the sites of insertion of the PGK-Neo-pCAG-hrGFP construct in the *CHD8*$^{+/GFP}$ clones cl. 1 and cl. 2. (**c**) *CHD8*$^{+/E1114;GFP}$ clones were generated by CRISPR/Cas9- and HDR-mediated insertion of point mutation (G to T) observed in the patients along a eGFP cassette into exon 16 of *CHD8*.(**d**) *CHD8*$^{+/S62X;GFP}$ cells were generated by CRISPR/Cas9- and HDR-mediated insertion of the point mutation (C to G) observed in the patients along a GFP cassette into exon 1 of *CHD8*. Southern blots confirmed corrected targeting in all cases **b-d** right panels.

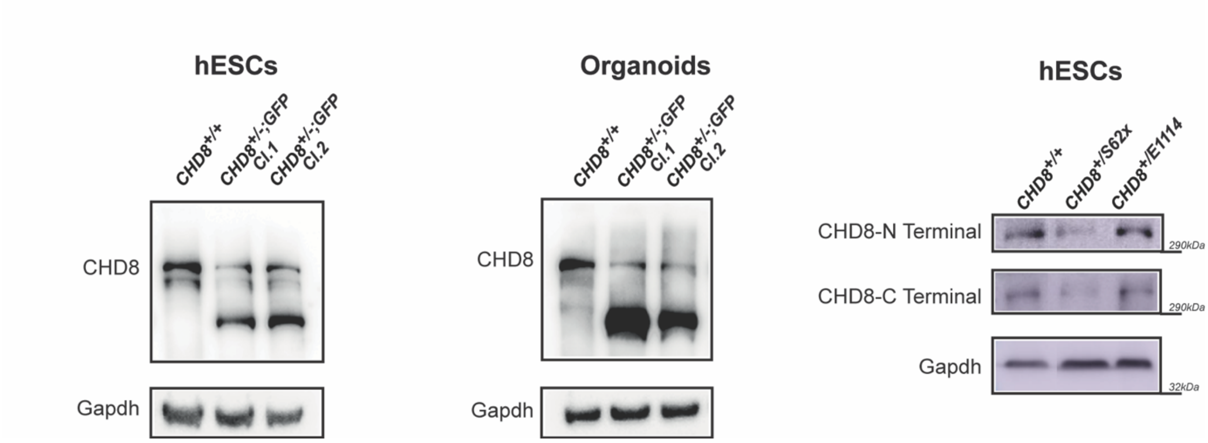


**Extended Data Figure 3. Western blot analysis of *CHD8* mutant hESC lines and cerebral organoids.** (**a**). Western blots for $CHD8^{+/+}$ and $CHD8^{+/-;GFP}$ (Cl. 1 and Cl. 2) in hESCs and cerebral organoids. (**b**) Western blots for $CHD8^{+/+}$, $CHD8^{+/S62X;GFP}$ and $CHD8^{+/E1114;GFP}$ in hESCs. Quantification revealed a reduction in CHD8 protein levels in mutant cells. Signal intensity was normalized to the wildtype for quantifications, GAPDH was used as loading control.

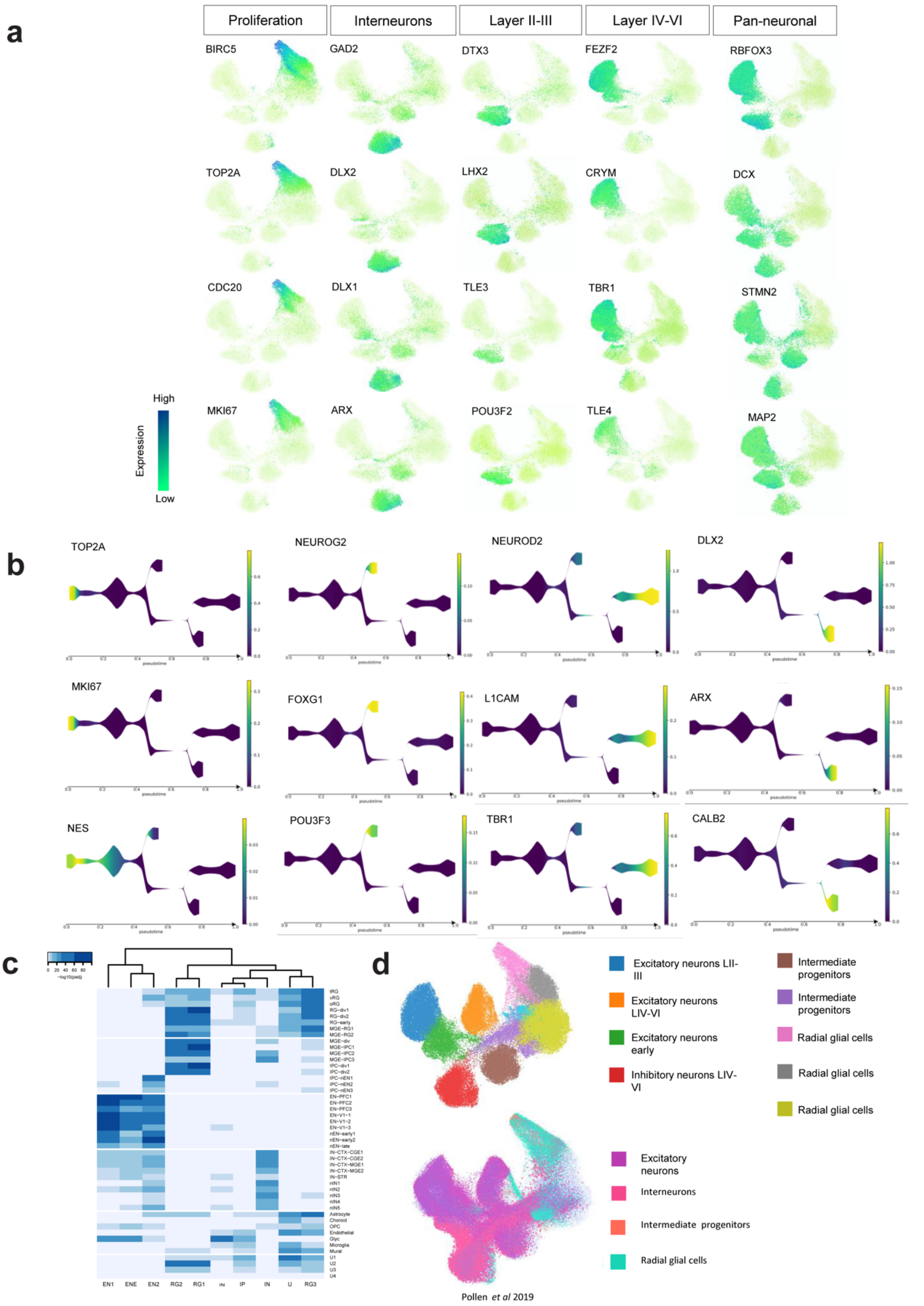


**Extended Data Figure 4. Cell populations and developmental trajectories are recapitulated in cerebral organoids.** (**a**) Representative gene expression profiles from different cellular identities and processes on the aggregated (stage/condition) dataset. (**b**) Tree-graph representation of gene expression patterns characteristic of different cell lineages. (**c**) Overlap of cluster markers between our clusters and an annotated human fetal brain single cell dataset[1]. (**d**) Projection of human fetal brain single cell dataset[2] on our generated cerebral organoid dataset (**d**) shows the overlap of pollen *et al* annotated cell-types, in the same projection.

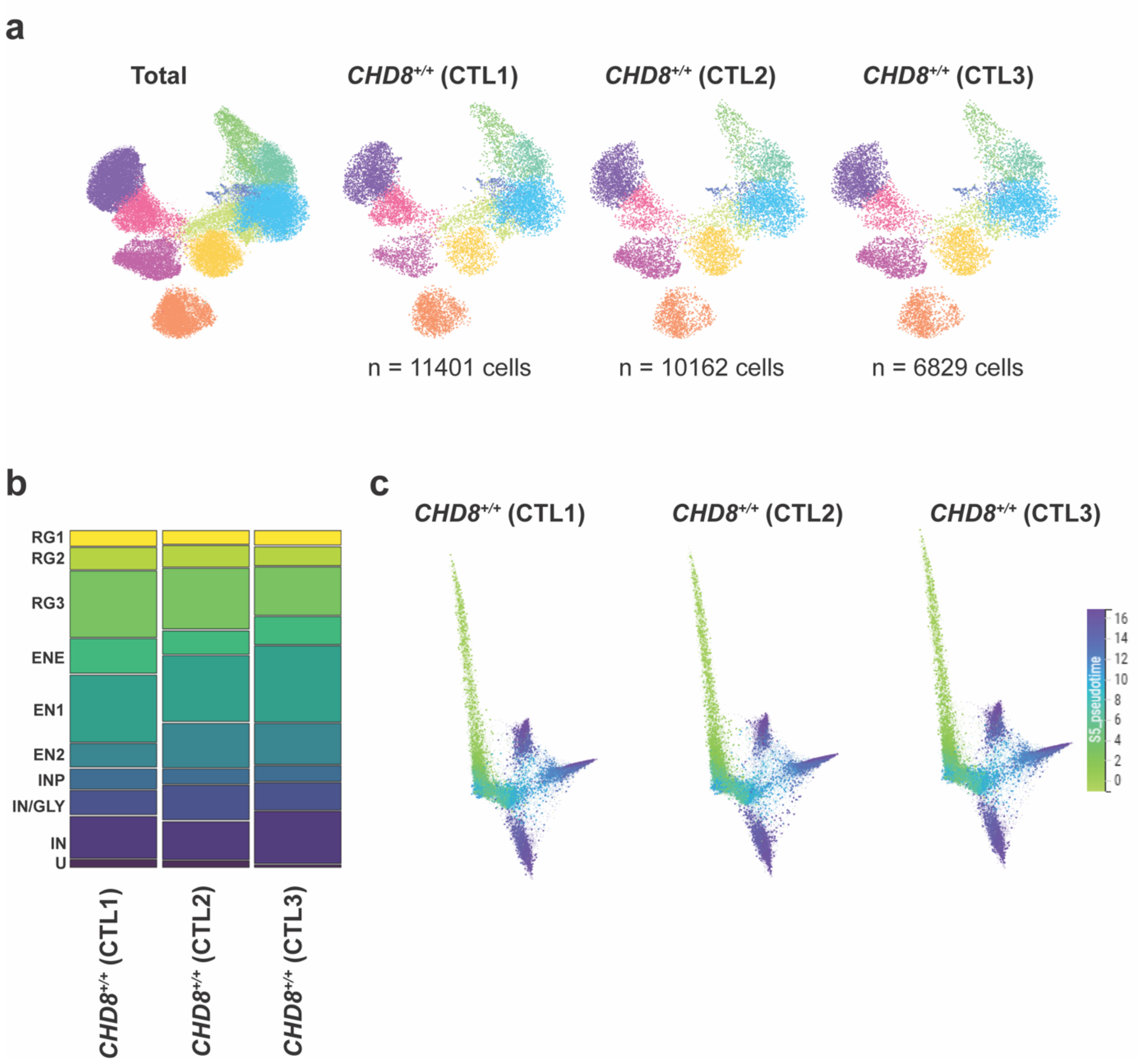


**Extended Data Figure 5. Cell type frequencies and differentiation trajectories are reproducible across lines.** (**a**) Distribution of cells across clusters visualized in UMAP of the three control lines. (**b**) Proportion of cells in each cluster on the aggregated stages across the 3 controls lines. (**c**) Diffusion map of control organoids depicting pseudotime trajectories in each individual line (origin:green to endpoint:blue).

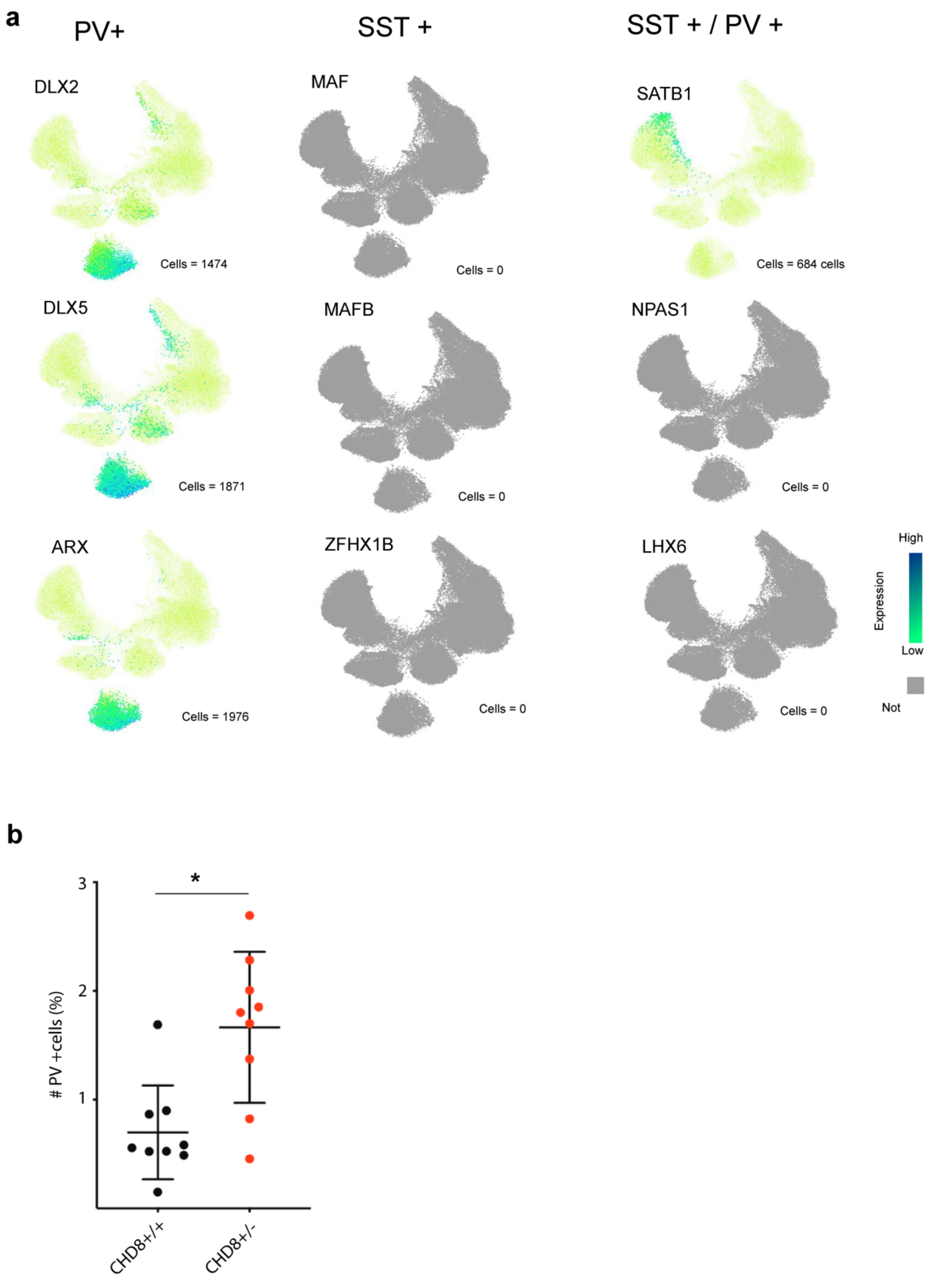


**Extended Data Figure 6. Characterization of developing interneuron identity.** (**a**) Number and distribution of cells expressing detectable levels of transcription factors characterizing the 3 main cortical interneuron identities[3]. (**b**) Quantification of PV+ positive cells at day 60 in control and mutant organoid samples. (Mann-Whitney test, *P < 0.01; *CHD8*$^{+/+}$, *n*= 9; *CHD8*$^{+/-}$, *n*= 9). N, number of nonconsecutive slices from at least 3 independent cerebral organoids.

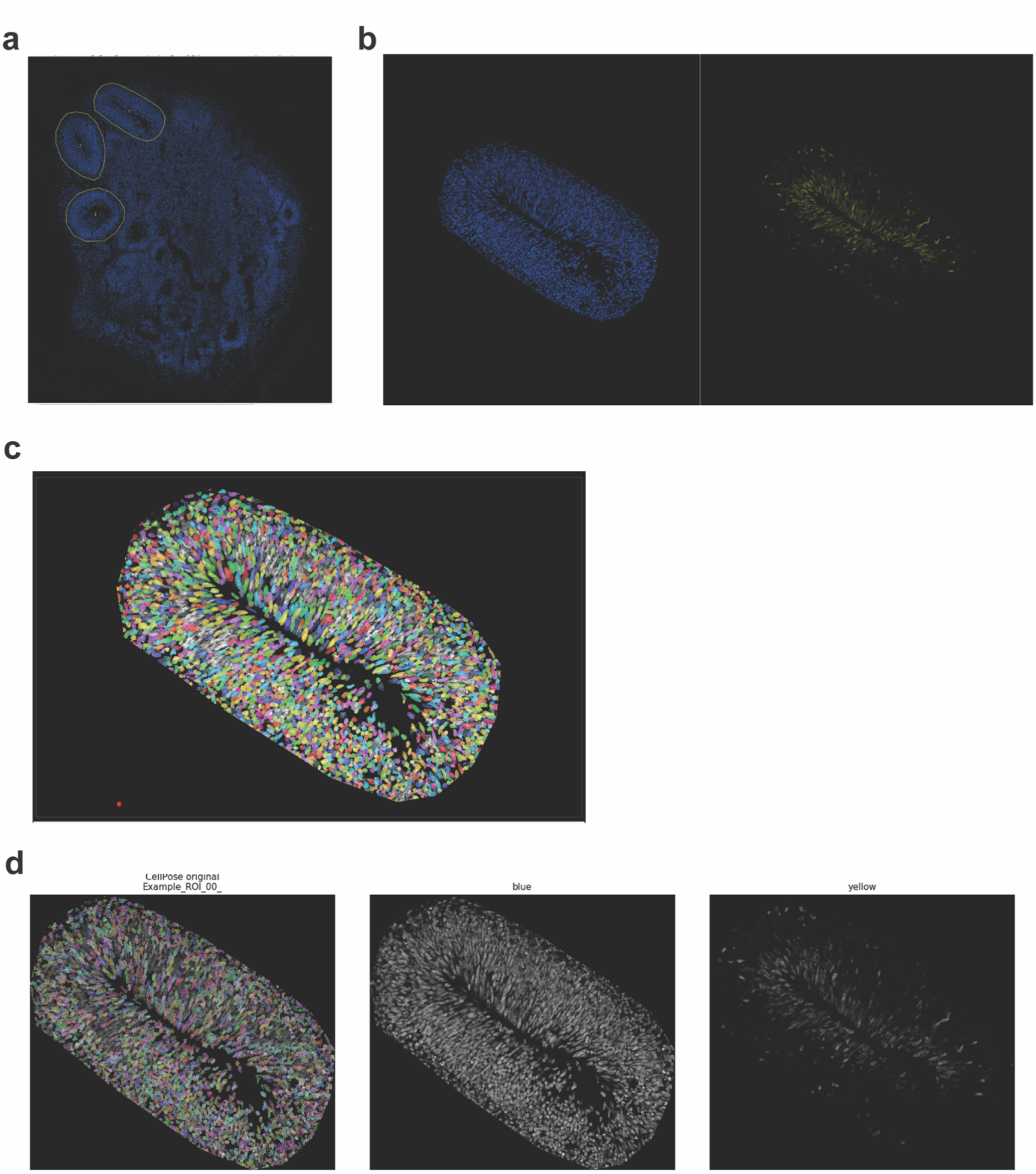


**Extended Data Figure 7. Semi-automatic cell counting pipeline. (a-b)** Representative images of rosettes selected for counting. **(c)** Masks for individual cells are computed with Cellpose. **(d)** The fitness of the masks is evaluated through a script conditional to predetermined parameters and provided to the end user for a manual last assessment.

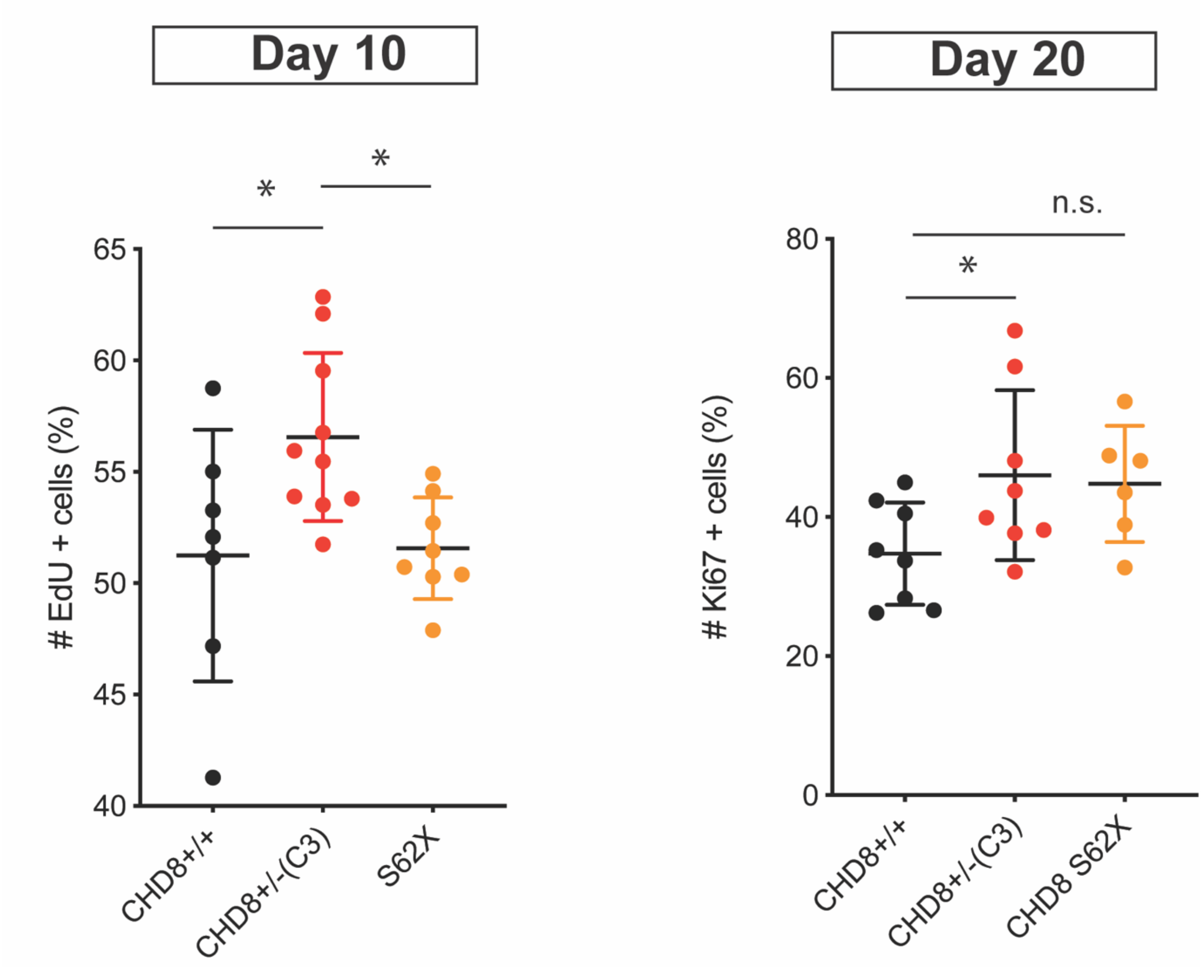


**Extended Data Figure 8. S62X mutation leads to a milder phenotype compared to *CHD8*$^{+/-}$ mutant organoids.** *CHD8*$^{+/S62X;GFP}$ mutant organoids show a non-significant trend in increased EdU+ cells at day 20 but not at day 10. (Day 10 *CHD8*$^{+/+}$, *n= 7; CHD8*$^{+/-}$, *n= 10; CHD8*$^{+S62;GFP-}$, *n= 8; day 20 CHD8*$^{+/+}$, *n= 8; CHD8*$^{+/-}$, *n= 8; CHD8*$^{+S62;GFP-}$, *n= 8).* N indicates number of organoids. *P ≤ 0.05, Ordinary One-way ANOVA followed by Dunnett's multiple comparisons test. Results are presented as mean ± s.d. Data for *CHD8*$^{+/+}$ and *CHD8*$^{+/-}$ presented also in Figure 4.

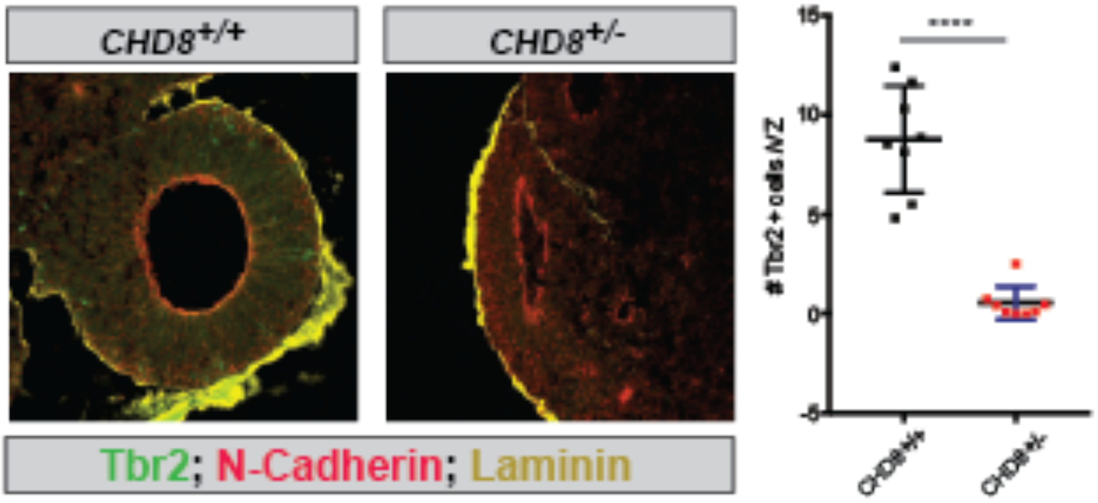


**Extended Data Figure 9. Decreased number of intermediate progenitors in *CHD8* mutant day 10 cerebral organoids.** Staining and quantification of Tbr2+ intermediate progenitors revealed a reduction of Tbr2+ cells in *CHD8*$^{+/-}$ (Unpaired t-test, P < 0.0001; *CHD8*$^{+/+}$, *n= 8; CHD8*$^{+/-}$, *n= 8*). ***P < 0.0001. N, number of cerebral organoids analyzed. Results are presented as mean ± s.d..


1 Nowakowski, T. J. *et al.* Spatiotemporal gene expression trajectories reveal developmental hierarchies of the human cortex. *Science* **358**, 1318-1323, doi:10.1126/science.aap8809 (2017).

2 Pollen, A. A. *et al.* Establishing Cerebral Organoids as Models of Human-Specific Brain Evolution. *Cell* **176**, 743-756 e717, doi:10.1016/j.cell.2019.01.017 (2019).

3 Lim, L., Mi, D., Llorca, A. & Marin, O. Development and Functional Diversification of Cortical Interneurons. *Neuron* **100**, 294-313, doi:10.1016/j.neuron.2018.10.009 (2018).
